# Multiscale mapping of venous remodelling in idiopathic pulmonary fibrosis

**DOI:** 10.64898/2026.09.17.752253

**Authors:** Daisuke Yamada, Huijuan Wang, Adam Szmul, Janik Riese, Ryoko Egashira, Coline H. M. van Moorsel, Nesrin Mogulkoc, Burak Gökçe, Shahab Aslani, Daryl Cheng, Yunjie Chen, Rahel Bodenmann, Matthias Brunner, Birger Tielemans, Joseph Brunet, Theresa Urban, Joanna Purzycka, Hector Dejea, Paul Tafforeau, Marie Vermant, Frouke T. van Beek, Britta Maurer, Sara Krättli, Marco Kreuzer, Remy Bruggmann, Tine Follet, Marcel Veltkamp, Tinne Goos, Kenny Anseele, Maarten Vander Kuylen, Hendrik W van Es, Klaus Engel, Recep Savas, Jose Manuel Brenes, Jeroen MH Hendriks, Thérése S Lapperre, Claire L Walsh, Danny Jonigk, Lavinia Neubert, Christopher Werlein, Hinrich Freitag, Sabine Dettmer, Jonas C Schupp, Laurens J de Sadeleer, Stephanie K. Robinson, Aurelie Fabre, Mark G. Jones, François Michel Carlier, Bart M Vanaudenaerde, Wim A Wuyts, Peter D Lee, Maximilian Ackermann, Stijn E Verleden, Janine Gote-Schniering, Joseph Jacob

## Abstract

The integrity of the pulmonary vasculature is a key determinant of lung health, yet challenges in visualisation and quantification have hindered interrogation of its pathophysiological role in chronic lung disease. Here, we align recent advances in microscale image acquisition with novel computer vision-based vessel segmentation models to demonstrate expansion of the bronchial and pulmonary veins across varying severities of tissue remodelling in idiopathic pulmonary fibrosis (IPF). Relating these imaging findings to molecular data in control, mild, and severe fibrosis, we show that bronchial venous endothelial cells expand beyond their physiological peribronchial niche even in mild disease, acquiring persistent angiogenic, inflammatory, and matrix-remodelling programmes that define a specialised fibrovascular-immune interface. Finally, we relate these microscale and molecular observations to clinical CT imaging, demonstrating that intrapulmonary vein enlargement independently associates with worsened survival across three IPF cohorts. Collectively, our multimodal, multiscale approach connects previously unresolved three-dimensional venous architecture to its molecular endothelial correlate, establishing venous enlargement as a prognostically significant and clinically relevant feature of IPF. Our analytical approach also demonstrates how biological discoveries made in intact ex vivo human organs can translate into measurable phenotypes in living patients, providing a template for other organs and diseases.

## INTRODUCTION

Idiopathic pulmonary fibrosis (IPF) is the most relentlessly progressive form of interstitial lung disease (ILD), yet the mechanisms driving irreversible fibrosis are incompletely understood^1^. Although epithelial injury and fibroblast activation dominate current models of disease pathogenesis, the accompanying reorganization of the pulmonary circulation, and its association with fibrotic progression, remain poorly characterized. This is despite growing evidence for the clinical and therapeutic relevance of the pulmonary vasculature^2^. Quantitative clinical CT studies have identified enlargement of intrapulmonary vessel-related structures as being one of the strongest imaging correlates of worsened patient outcome^3,4^. However, this quantitative measure combines arterial and venous compartments together with connected regions of fibrosis, leaving the question of whether a specific vascular compartment underlies this prognostic signal unresolved. Importantly, and distinct to the clinical scale, when considering the molecular and therapeutic perspectives, two of the internationally approved therapies for IPF are thought to act in part through antiangiogenic (nintedanib) and vasculoprotective (nerandomilast) mechanisms^5–7^.

The pulmonary venous system remains a particularly understudied component of the lung vasculature in pulmonary fibrosis. The pulmonary venous system comprises the bronchial veins that drain the systemic (bronchial arterial) circulation and the pulmonary veins that drain the pulmonary arterial inflow^8^. In practice however, these two venous circulations form a single interconnected venous pathway running through the peribronchovascular and subpleural connective tissues^9^. Delineating changes in the bronchial and pulmonary veins in IPF has been challenging, as their characterisation falls between the gaps of existing imaging and histological approaches. At the clinical scale, no tools reliably separate pulmonary arteries from veins on CT imaging of fibrotic lungs. At the microscopic scale, histopathological techniques preserve local tissue architecture but cannot trace venous networks continuously through the intact three-dimensional organ. At the molecular scale, single-cell and spatial profiling can resolve endothelial identity and molecular programmes, but only within very restricted tissue regions. Consequently, the three-dimensional architecture of venous remodelling in IPF has never been quantitatively mapped across the intact human lung, and no approach has connected molecular endothelial identity to organ-scale venous architecture and its clinically measurable counterpart.

Hierarchical phase-contrast X-ray tomography (HiP-CT) is an imaging modality that provides an opportunity to overcome this scale gap by enabling non-destructive three-dimensional imaging of intact human organs across multiple spatial scales while preserving vascular anatomy within their surrounding tissue architecture^10,11^. This new scale of image acquisition requires novel analytic segmentation approaches to convert a google earth-style view of an entire organ to a navigable google-maps-style demarcation of vascular pathways. Accordingly, we developed novel deep learning-based segmentations to define the venous system on whole-organ and zoom HiP-CT images and applied these to mild and severe regions of fibrosis.

Here, we provide a multiscale, multimodal approach that aligns three-dimensional HiP-CT imaging with novel computer vision-based vessel segmentation and molecular and spatial profiling to uncover venous remodelling across different fibrotic regions of IPF (**Fig. 1**). We reveal extensive three-dimensional expansion of the bronchial veins together with dilatation of higher-order pulmonary veins, already evident within mildly fibrotic lung regions. Integrated single-nucleus RNA sequencing, GeoMx spatial transcriptomics and multiplexed iterative immunofluorescence imaging resolved the endothelial correlate of this architectural remodelling, revealing expansion and reprogramming of bronchial venous endothelial cells within a specialised fibrovascular-immune interface. Finally, extending our computer vision-based segmentation approach to clinical CT imaging, we show that intrapulmonary venous enlargement independently associates with survival across three IPF cohorts. Together, our framework connects previously unresolved three-dimensional venous architecture to its molecular endothelial correlate and clinical manifestation, establishing venous remodelling as an early and prognostically significant feature of IPF.

**Figure 1.**
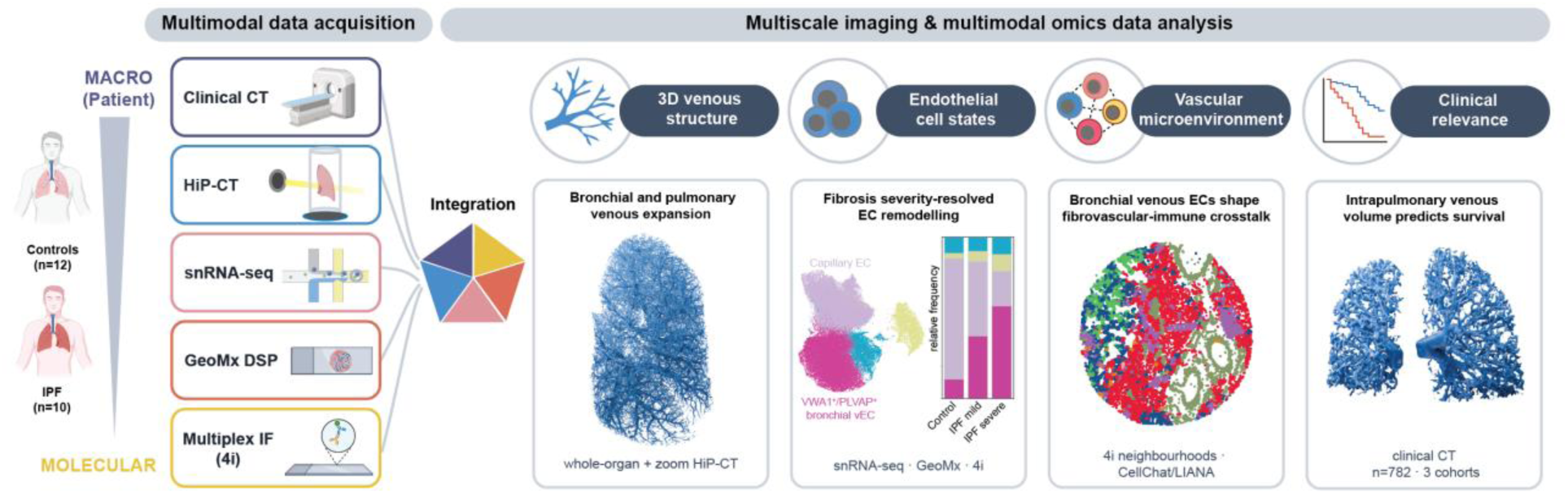
A multimodal, multiscale approach integrating imaging, molecular omics and clinical cohorts to resolve the venous system comparing healthy and IPF lungs. The study combines deep learning-based whole-lung venous segmentation and quantitative analysis, endothelial cell state characterisation, cellular crosstalk analysis and clinical outcomes to investigate vascular remodelling and its contribution to IPF pathogenesis. CT, computed tomography; HiP-CT, hierarchical phase-contrast tomography; snRNA-seq, single-nucleus RNA sequencing; GeoMx DSP, GeoMx digital spatial profiling; IF, immunofluorescence; 4i, iterative indirect immunofluorescence imaging; EC, endothelial cell; IPF, idiopathic pulmonary fibrosis; 3D, three-dimensional.

## RESULTS

### Whole-organ HiP-CT resolves three-dimensional venous remodelling across the fibrotic lung

To map vascular remodelling of an entire lung continuously across scales (**Fig. 2a**), we applied hierarchical phase-contrast tomography (HiP-CT) to lungs from patients with IPF from three centres in Belgium (Antwerp, Mont-Godinne and Brussels) and control lungs from Grenoble, France and Antwerp, Belgium (**Supplementary Table 1**). Whole lungs were imaged at isotropic voxel sizes of 16.6–42.3 µm and resampled to a common voxel size of 100 µm for whole-lung analysis. Whole organ imaging was used to resolve the pulmonary vein component (the bronchial circulation could not be resolved at the whole organ scale) of the pulmonary venous system (**Supplementary Movie 1**). At the whole lung scale, we developed a Forced Perspective segmentation approach which propagates sparse expert labels across imaging scales. The approach utilised iterative expert review and correction (**Fig. 2b**)^12^ to optimise segmentations of the pulmonary veins (PV) between training rounds. Based on the whole lung scan, areas containing mild and severe fibrosis were selected by expert radiologist readers (RY, DY, JJ) e. Zoom regions of interest were imaged at 2.5-5.1 µm and analysed after binning to 17-20 µm. The higher resolution of zoom imaging allowed visualisation of both the pulmonary veins and the bronchial veins in regions of fibrosis. At the zoom HiP-CT scale, the pulmonary and bronchial veins remain anatomically continuous and have no consistent boundary at which they can be divided. As a result, we used our Forced Perspective deep-learning pipeline to segment the pulmonary and bronchial veins as a single entity. Hereafter, all volumetric measurements of zoom images refer to the combined pulmonary venous system; where pulmonary and bronchial veins are described separately, this refers to qualitative evaluation of HiP-CT zoom imaging.

**Figure 2.**
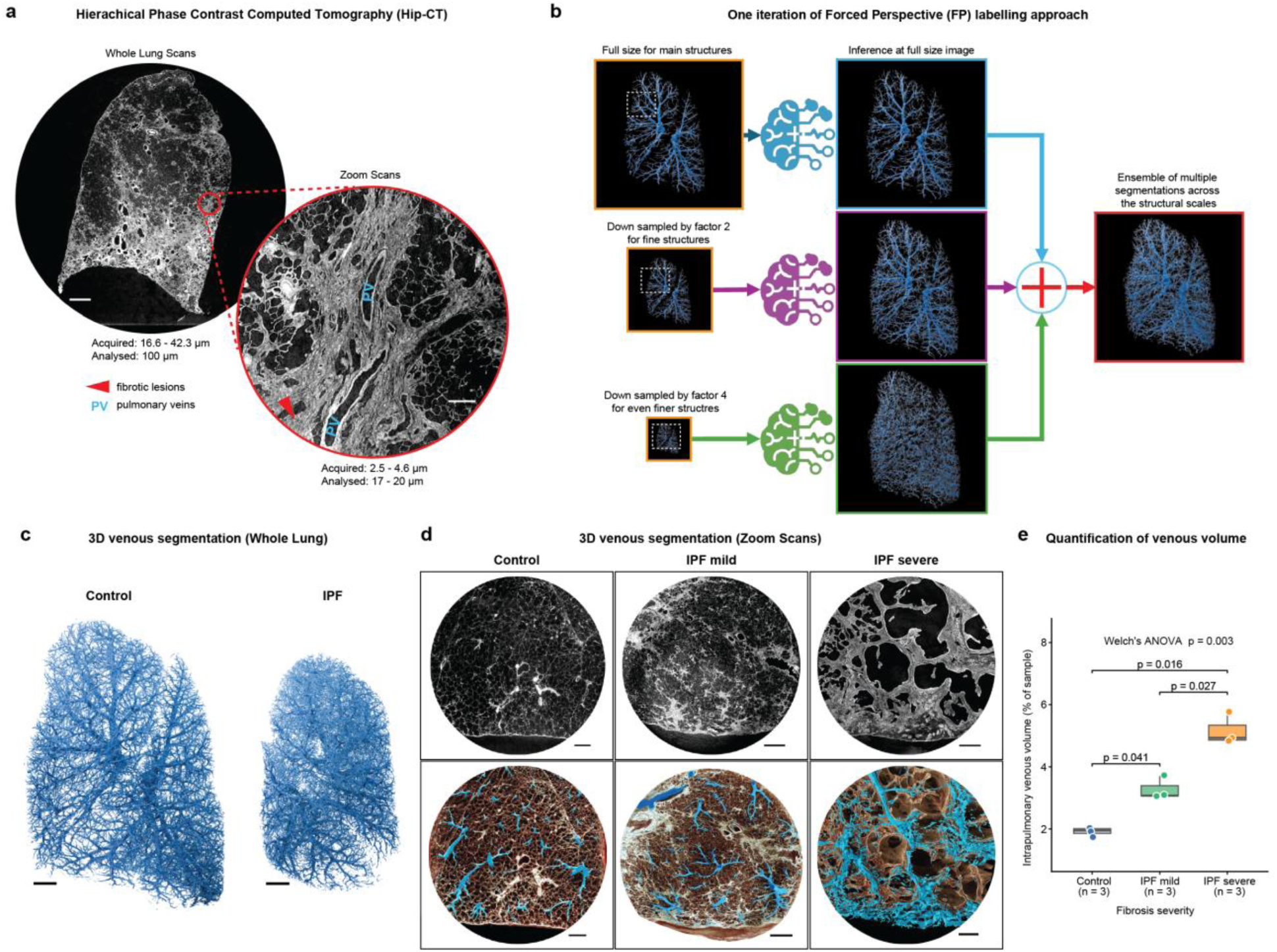
Whole-organ HiP-CT resolves three-dimensional venous remodelling across the fibrotic lung. Hierarchical phase contrast tomography scans of the whole lung, acquired at 16.6–42.3 µm isotropic voxel size and resampled to 100 µm for analysis (**a, left**), are shown with a corresponding zoom scan acquired at 2.5–5.1 µm and binned to 17–20 µm voxel size for analysis (a, right), where fibrotic tissue (red arrowheads) and pulmonary veins (PV) are highlighted. A novel deep learning approach was utilised using multiscale Forced Perspective (nnU-Net-based) segmentations to accurately segment the venous compartment at both imaging scales (**b**). The deep-learning segmentations created complex pulmonary vein trees at the whole lung level, where the bronchial circulation could not be resolved (**c**), and pulmonary and bronchial vein trees within zoom regions across severities of fibrotic remodelling in (**d**), the upper row shows the HiP-CT reconstructions and the lower row the corresponding venous segmentations. Pulmonary and bronchial veins were segmented and are rendered as a single venous compartment. Quantification of the combined pulmonary and bronchial venous volume (**e**) expressed as a percentage of total sample volume showed elevation of venous volume with increasing fibrosis severity (n = 3 volumes of interest per group; Welch’s one-way ANOVA, P = 0.003; pairwise Welch’s two-sample t-tests with Bonferroni correction, control versus mild P = 0.041, control versus severe P = 0.016, mild versus severe P = 0.027). Scale bars: 10 mm (a, whole lung), 750 µm (a, zoom), 15 mm (c), 3 mm (d).

Whole lung HiP-CT imaging revealed a distinct pattern of venous remodelling in IPF that differed by vessel order. Higher-order (more proximal) pulmonary veins draining regions of greater fibrotic severity were visibly dilated (**Fig. 2c**). In regions of mild fibrosis, where lung tissue retained a recognisable alveolar architecture, zoom images resolved large numbers of enlarged, tortuous bronchial veins with irregular, complex branching. Numerous anastomoses were evident between these vessels and proliferated, dilated pulmonary veins, with the latter draining into higher-order pulmonary veins (Fig. 2d). The size and number of bronchial and pulmonary veins increased in regions of more severe fibrosis. (**Fig. 2d**). In control tissue, visible bronchial veins were often solitary and did not form a distinct, comparable vascular network. The combined volume of pulmonary and bronchial veins was then calculated for each zoom HiP-CT volume of interest (control, mild, and severe IPF regions) and expressed as a percentage of total sample volume. Venous volume progressively increased with fibrosis severity (Fig. 2e), from a mean of 1.93 ± 0.15% in control tissue to 3.28 ± 0.37% in mild (P = 0.041) and 5.16 ± 0.52% in severe (P = 0.016 vs control, P = 0.027 vs mild) fibrotic regions.

### Progressive, lineage-specific endothelial remodelling in IPF resolved by multimodal single-cell and spatial profiling

HiP-CT imaging across control, mild and severe regions of fibrosis demonstrated extensive bronchial and pulmonary venous remodelling in areas of mildly fibrotic lung, with venous expansion becoming prolific in regions of severe fibrosis. To understand the cellular and molecular basis of pulmonary venous remodelling in IPF, we integrated three complementary modalities, single-nucleus RNA sequencing (snRNA-seq), GeoMx digital spatial profiling (DSP) and iterative indirect immunofluorescence imaging (4i), on regions of healthy lung tissue from control patients, and tissue regions in IPF patients stratified by microCT and histopathological assessment into mild and severe fibrosis (**Supplementary Figure 1**). Classification of fibrotic degree was confirmed by significant differences in microCT based alveolar surface density and histology-based Ashcroft score^13^ across the three groups (**Supplementary Figure 1 b,d**), enabling systematic characterisation of endothelial cell states, abundance and spatial organisation across lung tissue of increasing fibrotic severity.

snRNA-seq of 63,072 nuclei resolved 11,430 endothelial cells, annotated using a curated endothelial reference derived from the integrated Human Lung Cell Atlas^14^ together with the vascular endothelial taxonomy described by Schupp et al.^15^ (**Fig. 3a., Supplementary Figure 2a-g and 3a-h**). This resolved the major pulmonary endothelial populations, including aerocyte capillary (aCap; EDNRB, CA4), general capillary (gCap; FCN3, BTNL9), arterial (GJA5), lymphatic (LEC; PROX1, PDPN) and ACKR1⁺ venous endothelial cells, the latter comprising transcriptionally distinct pulmonary venous (CPE, CDH11) and bronchial venous populations (defined by VWA1 and PLVAP co-expression, hereafter referred to VWA1⁺/PLVAP⁺ bronchial venous EC (VWA1⁺/PLVAP⁺ bronchial vEC); **Fig. 3a, Supplementary Figure 3. b,f**). This last cell population has been described across previous studies using different nomenclatures, including systemic venous ECs^15^, (ectopic) bronchial ECs^16^, Bronch-1 ECs^17^, and ectopic COL15A1⁺ pVECs^18^. CIBERSORTx-based deconvolution of 42 CD31⁺ endothelial-rich GeoMx DSP regions of interest recovered the same endothelial populations across spatially heterogeneous fibrotic niches (**Supplementary Figure 4 and 5**)^19–21^. Independent 4i imaging using a 24-marker antibody panel resolved the corresponding broader endothelial compartments at the protein level, including capillary (PRX⁺), lymphatic (PDPN⁺) and VWA1⁺/PLVAP⁺ bronchial vEC, with the remaining CD31⁺ endothelium grouped as an “Other CD31^+^” cell population (**Fig. 3b, Supplementary Figure 6**)^22^.

**Figure 3.**
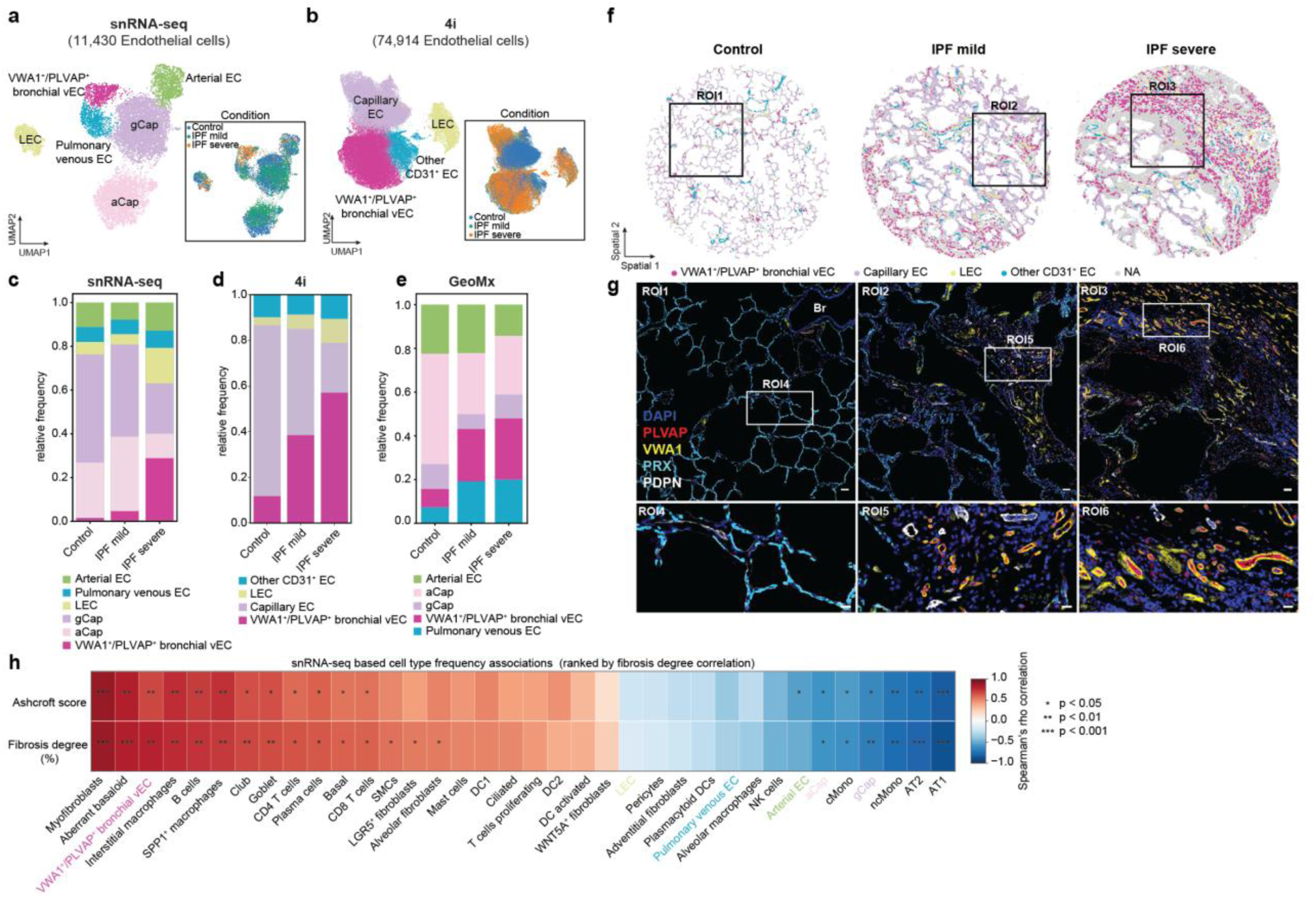
Multimodal analyses consistently reveal the expansion of VWA1⁺/PLVAP⁺ bronchial venous EC in IPF patients. UMAP embedding of endothelial cells from (a) snRNA-seq (11,430 cells) and (b) 4i, colored by endothelial cell subtypes and disease condition. Relative endothelial cell composition across (c) snRNA-seq, (d) 4i, and (e) GeoMx datasets demonstrating a consistent expansion of VWA1⁺/PLVAP⁺ bronchial venous EC in IPF. (f) Representative spatial maps of endothelial cell subtypes in control, IPF mild and IPF severe human lung tissues. (g) Higher magnification 4i images showing the VWA1⁺/PLVAP⁺ bronchial venous EC (red and yellow), PRX (blue, capillary) and PDPN (white, LEC) from control, IPF mild and IPF severe. VWA1⁺/PLVAP⁺ bronchial venous EC restricted to the peribronchial region in control lungs (ROI4) but extended beyond this region in IPF mild (ROI5) and IPF severe (ROI6) lung tissues. (scale bar: ROI 1-3, 50 µm; scale bar: ROI4-6, 20 µm). (h) Spearman correlation with FDR correction between the snRNA-seq-based relative cell type frequency and Ashcroft score or fibrosis degree (in percent) of the individual tissue samples. The cell types are ranked according to their correlation with fibrosis degree. Biological replicates: snRNA-seq: n=7 control, n=6 IPF patients, GeoMx/4i: n=4 control, n=8 IPF.

Across fibrosis stages, all three modalities revealed a consistent, lineage-specific reorganisation of the endothelial compartment, characterised by progressive capillary depletion and selective expansion of the bronchial venous lineage (**Fig. 3c–e, Supplementary Figures 2h-n, 5a-c, 7a-e**). In control lungs, aCap and gCap endothelial cells dominated the endothelial compartment, with 4i spatially localising PRX⁺ capillary endothelial cells throughout the alveolar microvasculature, whereas VWA1⁺/PLVAP⁺ bronchial vEC were comparatively sparse and largely confined to their expected peribronchial distribution (**Fig. 3f–g; ROI1**). Already in mild fibrosis, capillary endothelial abundance declined while VWA1⁺/PLVAP⁺ bronchial vEC increased and extended beyond their peribronchial niche into remodelled alveolar parenchyma, paralleling the early structural venous remodelling identified by HiP-CT. With increasing fibrosis severity, this divergence became more pronounced, with further loss of aCap and gCap populations and progressive expansion of VWA1⁺/PLVAP⁺ bronchial vEC, which became the predominant endothelial population in advanced fibrosis and occupied extensive remodelled parenchymal regions (**Fig. 3f–g; ROI2– ROI6**), in keeping with the ectopic redistribution of systemic/bronchial venous endothelium described in fibrotic human lung^15,16,23,24^. Finally, correlation of snRNA-seq-derived cell-type frequencies with histological fibrosis severity (Ashcroft score and tissue fibrosis extent) identified VWA1⁺/PLVAP⁺ bronchial vEC as the endothelial population most strongly associated with fibrosis, ranking alongside canonical fibrogenic cell states including myofibroblasts and aberrant basaloid epithelial cells (**Fig. 3h**). This association was independently reproduced in the 4i dataset (**Supplementary Figure 7f)** and supported by a concordant, although non-significant, trend in GeoMx DSP (**Supplementary Figure 5d**). By contrast, capillary (aCap and gCap), arterial and pulmonary venous EC were negatively associated with fibrosis, suggesting that venous remodelling in IPF mainly involves the bronchial venous lineage.

### Distinct stage-specific endothelial programs define capillary and bronchial venous remodelling in IPF

To identify the endothelial populations undergoing the greatest transcriptional remodelling during IPF, we applied Augur perturbation analysis^25^ to the snRNA-seq dataset. Consistent with previous reports of endothelial dysfunction in human IPF, capillary and venous endothelial populations ranked among the most perturbed cellular lineages relative to control (**Fig. 4a, Supplementary Figure 8g**). Notably, both compartments were already transcriptionally perturbed in mild fibrosis, indicating that molecular remodelling accompanies the early structural vascular changes identified by HiP-CT.

**Figure 4.**
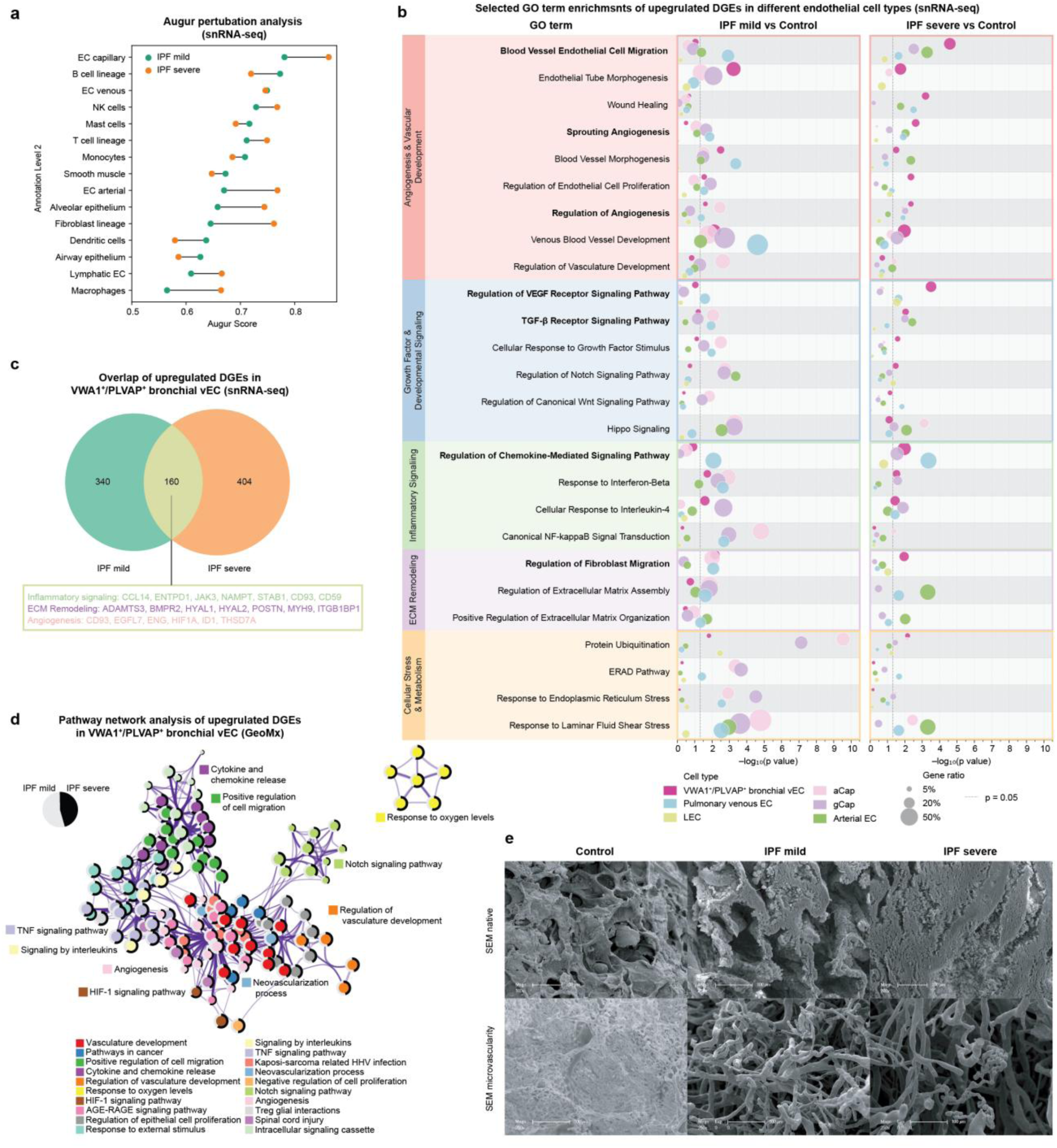
Pathway analyses reveal angiogenesis and vascular remodelling in IPF lungs. (a) Augur perturbation analysis of snRNA-seq datasets prioritizing broad cell populations (level_2 annotation) according to their transcriptional perturbation comparing IPF mild vs control and IPF severe vs control. (b) Selected GO terms from pathway overrepresentation analysis of upregulated genes in different endothelial cell types comparing IPF mild vs Control, IPF severe vs control (log2FC > 0.3, adjusted P value <0.2). Selected GO terms are grouped into distinct biological theme labelled with different colors. The dashed line indicates raw P value of 0.05. (c) Venn diagram showing the overlapping upregulated genes in VWA1⁺/PLVAP⁺ bronchial venous EC from IPF mild vs control and IPF severe vs control from the snRNA-seq dataset. Representative overlapping genes are highlighted and grouped into different biological categories with different colors. (d) Pathway network analysis of upregulated genes in VWA1⁺/PLVAP⁺ bronchial venous EC from GeoMx datasets. (e) Scanning electron microscopy (SEM) images of an age-matched control lung, and regions of mild and severe lung fibrosis. The images in the top row show the native SEM images coated with a conductive layer of gold. The bottom row demonstrates corrosion casting of the lung microvasculature after tissue clearing. In the control sample alveoli are visible (top left image) and there is a fine lattice-like capillary branching network visible. In a region of mild fibrosis, fibrotic remodelling is evident with somewhat preserved tissue architecture (top, middle), but the capillary network is destroyed with irregular sprouting and branching of larger vessels already evident (bottom, middle). In a region of severe fibrosis, there is extensive fibrotic remodelling of tissue (top right) and loss of vasculature with only limited numbers of abnormally branching arterioles/venules visible (bottom right).

Gene ontology enrichment of differentially expressed genes (log_2_FC>0.3, adjusted P value<0.2) revealed lineage-specific transcriptional programmes that differed according to endothelial subtype and fibrosis stage (**Fig. 4b**; **Supplementary Files 1 and 2, Supplementary Figure 8c-h).** Capillary endothelial cells (aCap and gCap) preferentially activated cellular stress and adaptive programmes, including protein ubiquitination, endoplasmic reticulum (ER) stress and ER-associated degradation (ERAD), together with inflammatory signalling through interferon-β and canonical NF-κB pathways. Developmental signalling through Notch and Hippo pathways was also enriched, consistent with activation of endothelial injury-response and metabolically stressed programmes previously found in experimental and aged fibrotic lung^24,26^. In parallel, VWA1⁺/PLVAP⁺ bronchial vEC exhibited enrichment of angiogenic and vascular-remodelling pathways, including sprouting angiogenesis, blood vessel endothelial cell migration, regulation of angiogenesis and VEGF and TGF-β receptor signalling. These programmes were already detectable in mild fibrosis and became progressively more prominent with increasing fibrosis severity, accompanied by enhanced chemokine-mediated signalling together with fibroblast migration pathways (**Fig. 4b)**. This progressive activation of angiogenic and matrix-remodelling programmes suggested that VWA1⁺/PLVAP⁺ bronchial vEC may represent an active signalling interface between vascular and stromal compartments during fibrosis progression.

To determine whether this phenotype was maintained throughout disease progression, we compared genes upregulated in VWA1⁺/PLVAP⁺ bronchial vEC between mild and severe fibrosis. Despite progressive transcriptional remodelling, 160 genes were shared between disease stages (**Fig. 4c**), demonstrating that a substantial component of the bronchial venous programme is established early and retained throughout disease progression. This conserved signature included inflammatory mediators (e.g. CCL14, CD59), extracellular matrix-remodelling genes (e.g. POSTN) and angiogenesis-associated regulators (e.g. HIF1A), consistent with coordinated inflammatory activation, vascular remodelling and matrix deposition.

Spatial validation by GeoMx DSP confirmed that these transcriptional programmes were maintained within the spatially resolved tissue microenvironment. In particular, the spatially preserved enrichment of angiogenic, hypoxia-associated and matrix-remodelling modules supported the emergence of a specialised vascular microenvironment rather than a generalized endothelial stress response. Pathway network analysis of CIBERSORTx-deconvolved VWA1⁺/PLVAP⁺ bronchial vEC transcriptomes independently reproduced interconnected modules related to angiogenesis, HIF-1 signalling, cytokine and chemokine signalling, positive regulation of cell migration and hypoxia responses (**Fig. 4d, Supplementary Figure 9a,b),** demonstrating that the angiogenic and inflammatory phenotype identified by dissociated snRNA-seq is spatially preserved within the vascular compartment of fibrotic lesions.

Finally, scanning electron microscopy of vascular corrosion casts provided ultrastructural correlates of both endothelial programmes (Fig. 4e). Compared with the fine, lattice-like alveolar capillary network in control lungs, regions of mild fibrosis showed disruption of the capillary network with irregular sprouting and branching of larger vessels, consistent with active angiogenic remodelling, whereas regions of severe fibrosis showed extensive loss of the capillary bed, with only a few abnormally branching arterioles and venules remaining.

Together, these orthogonal analyses demonstrate that capillary and VWA1⁺/PLVAP⁺ bronchial vEC mount distinct but concurrent responses with increasing fibrosis severity. Whereas capillary endothelial cells predominantly activate stress-associated programmes, VWA1⁺/PLVAP⁺ bronchial vEC establish an angiogenic, inflammatory and matrix-remodelling programme that is already evident in mild fibrosis and persists as this compartment further expands with increasing fibrosis severity.

### VWA1⁺/PLVAP⁺ bronchial venous EC define a fibrovascular interface surrounded by lymphoid cells in fibrotic lung tissue

To determine how VWA1⁺/PLVAP⁺ bronchial vEC are organised within the fibrotic microenvironment, we interrogated their local cellular context using the single-cell 4i dataset. Unsupervised spatial neighbourhood analysis identified reproducible cellular niches (CNs) based on local cell composition, one of which represented a distinct fibrovascular niche enriched for VWA1⁺/PLVAP⁺ bronchial vEC together with lymphatic endothelial cells and fibroblasts, including both POSTN⁺ activated fibroblasts and other CD90⁺ fibroblast populations, yet not POSTN^+^/TNC^+^ myofibroblasts which formed a separate niche together with aberrant basaloids, club/goblet and basal cells^27^ (**Fig. 5a, Supplementary Figure 10a/b**). Spatial mapping demonstrated that these cell types consistently co-localised within shared tissue territories rather than distributing randomly throughout the parenchyma (**Fig. 5b-d, Supplementary Figure 10a/b**). Quantification across disease stages showed progressive expansion of this fibrovascular neighbourhood from control through mild to severe fibrosis (**Fig. 5e**), indicating that its establishment accompanies progressive tissue remodelling.

**Figure 5.**
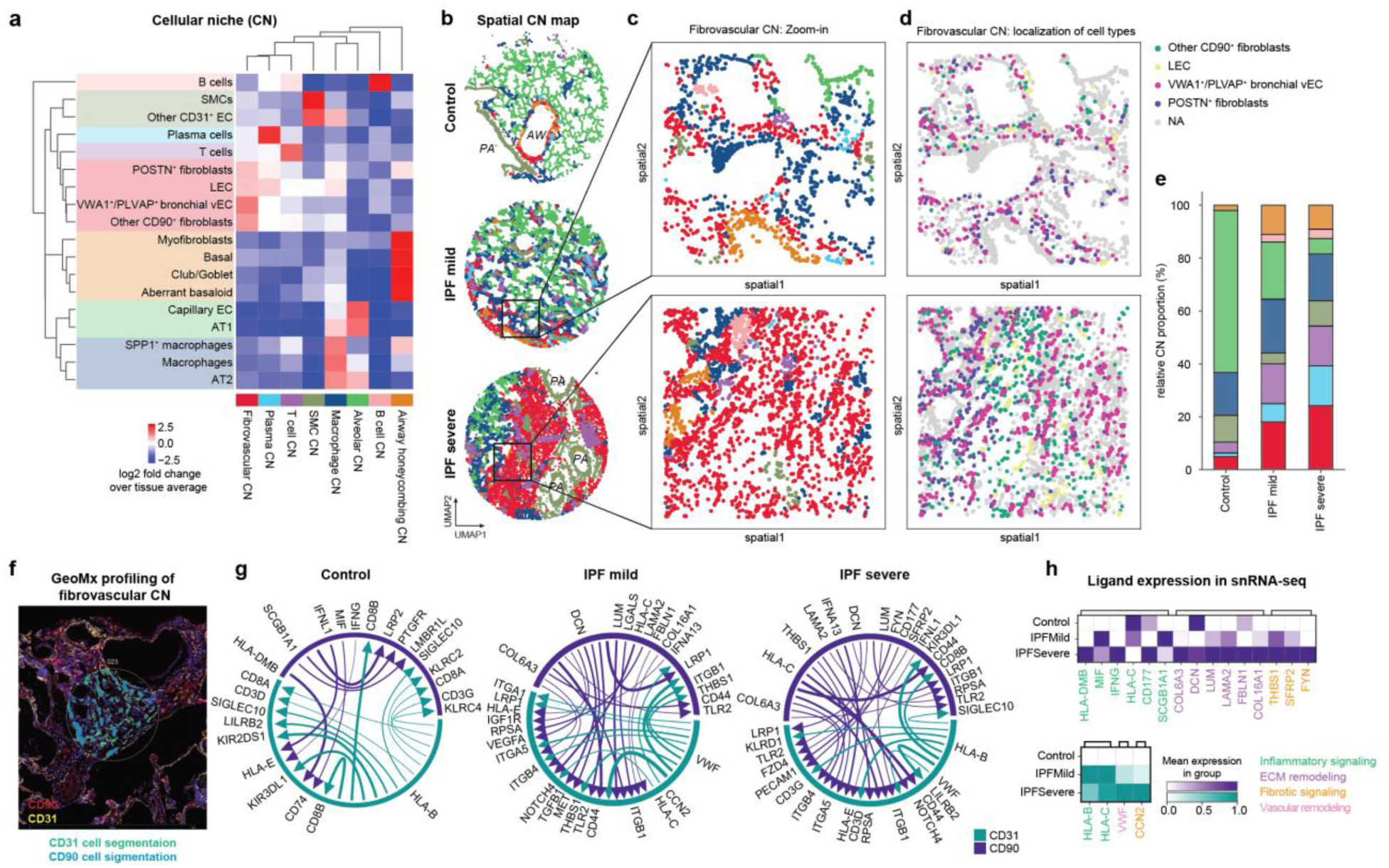
Spatial profiling by 4i and GeoMx identifies a fibrovascular niche enriched in IPF. (a) Spatial neighborhood analysis of 18 cell populations identified by 4i, grouped into 8 distinct cellular niches (CNs). (b) Representative spatial CN maps in control, IPF mild and IPF severe. (c) Higher magnification images highlighting the identified fibrovascular CN in IPF mild and IPF severe lung tissues. (d) Corresponding spatial image showing cellular composition of the fibrovascular niche. (e) Relative proportion of each cellular niche across disease severity. (f) Representative GeoMx spatial profiling of the fibrovascular niche, white circle demarcates area in which segmentation of the CD31^-^ and CD90^-^positive compartment was performed. (g) Predicted ligand-receptor interactions between CD31^+^ endothelial (green) and CD90^+^ fibroblast (purple) populations within GeoMx profiles from control, IPF mild and IPF severe regions. Arrow thickness is corresponding to interaction strength. (h) Expression of GeoMx-identified ligands involved in endothelial-fibroblast communication in CD31^+^ endothelial (green) and CD90^+^ fibroblast (purple) populations in the snRNA-seq dataset. Ligands are classified into 4 biological categories highlighted by different colors.

To investigate communication underlying the expanding fibrovascular niche, we performed spatial ligand–receptor analysis between adjacent CD31⁺ endothelial and CD90⁺ stromal regions using GeoMx DSP^28^ (**Fig. 5f, Supplementary Figure 4d**). Consistent with the progressive expansion of the bronchial venous endothelial compartment, ligand–receptor interactions increased in both number and complexity with increasing fibrosis severity, indicating progressive strengthening of endothelial–stromal crosstalk within the niche (**Fig. 5g**). In control lungs, signalling was primarily characterised by MHC class I and II interactions (HLA–CD74, HLA–CD8, HLA–KIR), consistent with physiological antigen presentation and immune homeostasis rather than active tissue remodelling. By contrast, regions with mild fibrosis showed a marked shift towards extracellular matrix-dependent communication. CD90⁺ stromal regions displayed prominent collagen (COL6A3, COL16A1), laminin (LAMA2), decorin (DCN), lumican (LUM) and fibulin (FBLN1) signalling converging on integrins (ITGB1, ITGA1, ITGA5), LRP1 and CD44, indicative of active matrix remodelling and mechanotransduction. Endothelial regions simultaneously exhibited increased VWF- and CCN2-associated signalling together with interactions involving VEGFA, TGF-β, IGF1R and NOTCH4 receptors, consistent with endothelial activation, angiogenic remodelling and profibrotic signalling. In severe fibrosis regions, these matrix-dependent interactions became further reinforced and were accompanied by the emergence of immune-regulatory signalling. Collagen-, laminin- and proteoglycan-derived ligands (COL6A3, LAMA2, DCN and LUM) remained dominant, together with increased THBS1 signalling, reflecting a mature fibrotic extracellular matrix. Notably, severe fibrosis also demonstrated increased engagement of inhibitory immune receptors, including KIR3DL1, SIGLEC10 and LILRB2, together with MHC class I ligands, suggesting establishment of an immunoregulatory vascular interface capable of coordinating lymphoid cell interactions.

To independently corroborate these spatial interactions, we examined the average expression of the corresponding ligands within the endothelial (CD31⁺) and fibroblast (CD90⁺) compartments in the snRNA-seq dataset (**Fig. 5h**). Consistent with the GeoMx analysis, expression of multiple inflammatory, angiogenic, extracellular matrix-associated and profibrotic ligands increased with fibrosis severity in both compartments. In particular, ligands associated with collagen- and laminin-mediated matrix signalling, endothelial activation and immune regulation recapitulated the stage-dependent communication identified by spatial ligand–receptor profiling

Several ligands detected within the CD31⁺ and CD90⁺ GeoMx regions were not expressed by the corresponding endothelial or fibroblast compartments in the dissociated snRNA-seq dataset, and both ROI types contained lymphoid-associated transcripts, including CD8A/B or CD3G. Together with the emergence of immune-regulatory ligand–receptor interactions in severe fibrosis, this indicated that the spatially profiled regions captured transcripts from cells immediately adjacent to the fibrovascular niche. To define this surrounding microenvironment, we returned to the 4i dataset and performed patch proximity analysis, quantifying the cell types enriched around the fibrovascular niche across fibrosis stages (**Fig. 6a, Supplementary Figure 10 c-e**). This revealed a progressive, stage-dependent accumulation of lymphoid cells, with T cells, B cells and plasma cells increasingly represented around the fibrovascular CN in mild and severe fibrosis relative to control (**Fig. 6a,b**). These populations were concentrated around the niche rather than distributed evenly through the parenchyma, forming dense peri-niche immune aggregates in severe disease (**Fig. 6b**), and high-dimensional 4i imaging validated this organisation in situ, showing VWA1⁺/PLVAP⁺ bronchial vECs surrounded by CD3⁺ T cells, CD20⁺ B cells and MZB1⁺ plasma cells within remodelled fibrotic regions (**Fig. 6c**). This organisation in severe fibrosis is consistent with the immune niche of lymphoid foci surrounded by remodelled endothelial vessels described in end-stage IPF^29^, which our stage-resolved, lineage-resolved analysis localises to the bronchial venous compartment and identifies already in mild fibrosis.

**Figure 6.**
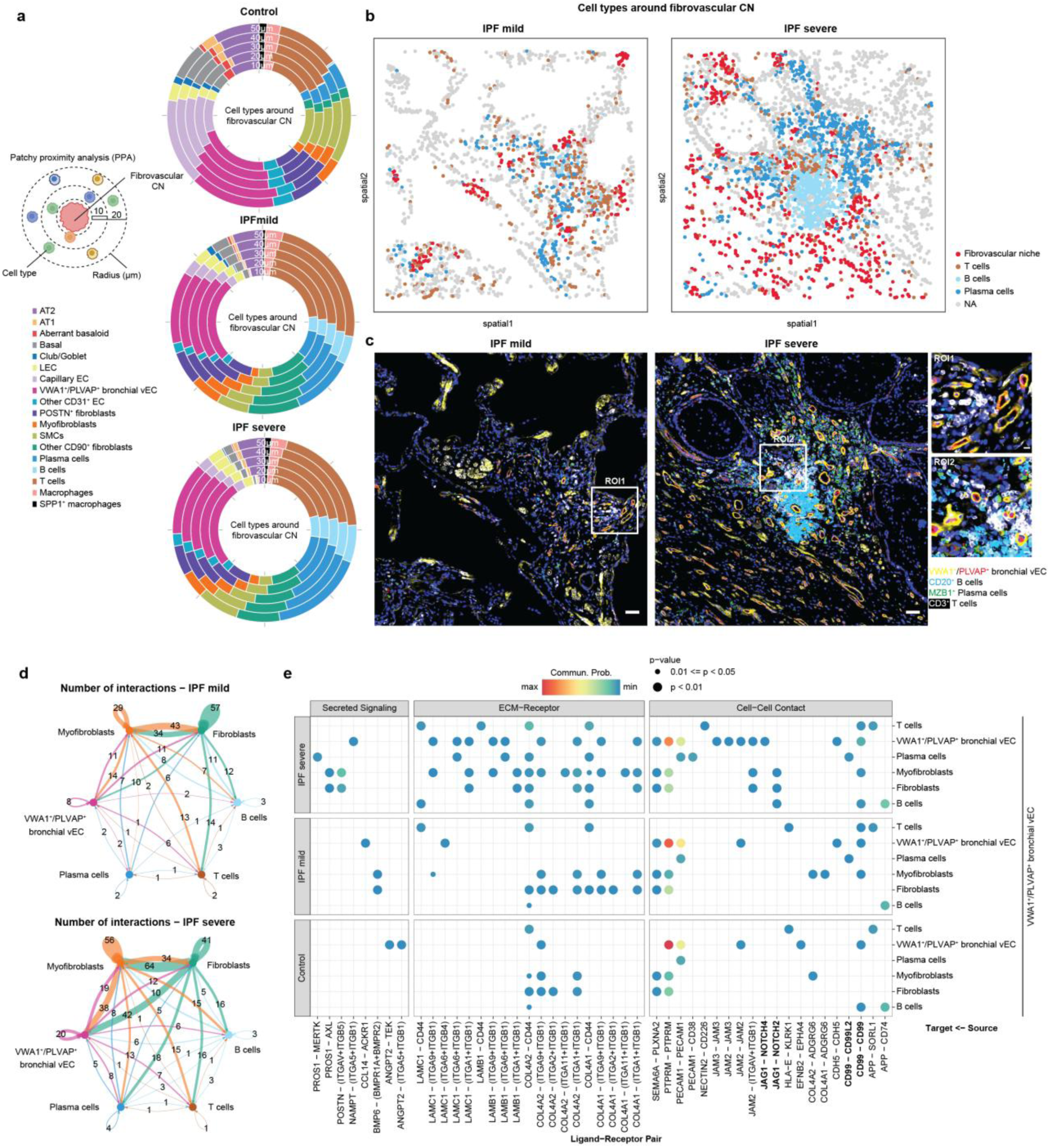
Spatial analyses reveal increased immune cell accumulation around fibrovascular niche in IPF. (a) Schematic illustration of the patchy proximity analysis (left). PPA showing the cellular composition surrounding the fibrovascular niche within radii of 10, 20, 30, 40 and 50 µm in control, IPF mild and IPF severe lungs (right). (b) Representative spatial images highlighting the enrichment of B cells, T cells and plasma cells surrounding the fibrovascular niche in IPF mild and severe patients. (c) Corresponding 4i images validating the spatial localization of CD20^+^ B cells (light blue), MZB1^+^ plasma cells (green) and CD3^+^ T cells (white) adjacent to VWA1⁺/PLVAP⁺ bronchial venous EC (red and yellow). Scale bar, 50 µm (overview) and 10 µm (magnified views). (d) Circle plots showing inferred cell-cell communications among VWA1⁺/PLVAP⁺ bronchial venous EC, fibroblasts and neighbouring immune cell populations in IPF mild and IPF severe lungs by CellChat analysis in the snRNA-seq dataset. (e) Bubble plot summarizing the CellChat-derived ligand-receptor interactions originating from VWA1⁺/PLVAP⁺ bronchial venous EC and targeting VWA1⁺/PLVAP⁺ bronchial venous EC, fibroblasts and neighbouring immune cell populations across control, IPF mild and IPF severe groups. Interactions with P > 0.05 were excluded. Dot size represents the level of statistical significance, with larger dots indicating lower P values. Dot color represents estimated communication probability, ranging from lower (blue) to higher (red) probability.

To characterise the signalling underlying this multicellular organisation, we performed CellChat analysis of VWA1⁺/PLVAP⁺ bronchial vEC and their neighbouring immune and stromal populations^30^ (**Fig. 6d,e**). The number of inferred interactions increased markedly from mild to severe fibrosis, indicating progressive intensification of crosstalk within the niche (**Fig. 6d, Supplementary Figure 10f**). Consistent with the GeoMx ligand–receptor analysis, VWA1⁺/PLVAP⁺ bronchial vEC engaged fibroblasts, myofibroblasts and lymphoid populations through secreted, ECM-receptor and cell-cell contact pathways (**Fig. 6e, Supplementary Figure 11**). Secreted and ECM-receptor signalling was dominated by laminin and collagen interactions engaging integrin receptors, recapitulating the matrix-dependent endothelial–stromal communication identified spatially, whereas cell-cell contact signalling included adhesion and immune-regulatory pathways (e.g. CD99, MHC class I) linking endothelial cells to lymphoid populations, consistent with the emergence of immune-regulatory signalling observed in severe fibrosis by GeoMx. Together, these analyses define a multicellular fibrovascular microenvironment in which endothelial, fibroblast and immune compartments physically and molecularly converge around the bronchial venous vasculature, with this convergence intensifying as disease severity increases.

### Clinical CT reveals venous expansion with prognostic significance in IPF

Having established that venous remodelling is an early feature of IPF and is embedded within a fibrovascular-immune niche, we next asked whether this vascular phenotype could be resolved on routine clinical imaging and could provide prognostic information in patients with IPF. We therefore translated the venous enlargement observed by zoom and whole-lung HiP-CT to clinical-scale CT imaging in patients with IPF and examined its association with survival.

We analysed clinical CT imaging in three independent IPF cohorts (Ege University Hospital, Izmir, Türkiye, n = 298; St Antonius Hospital, Nieuwegein, Netherlands, n = 239; UZ Leuven, Leuven, Belgium, n = 245; total n = 782 patients and CTs; **Fig. 7a, Supplementary Table 2, Supplementary Figure 12**). Segmentation pipelines and expertise developed from analysis of whole-lung HiP-CT imaging were used to inform analysis of the intrapulmonary veins on clinical CT imaging. The Forced Perspective methodology was again employed, with iterative human-in-the-loop corrections of segmentation masks as detailed in Methods. Intrapulmonary venous volume was expressed as a percentage of total lung volume (%PV; **Fig. 7b-e**). The three IPF cohorts differed in several respects: the Belgian cohort was older and universally treated with antifibrotic therapy, the Dutch cohort had the highest proportion of ever-smokers and the lowest antifibrotic exposure, and the Turkish cohort had the lowest baseline FVC. The cohorts primarily comprised early to moderate stage disease (GAP stage I = 47.1%, GAP stage II = 43.7%, versus GAP stage III = 9.2%; Supplementary Table 2).

**Figure 7.**
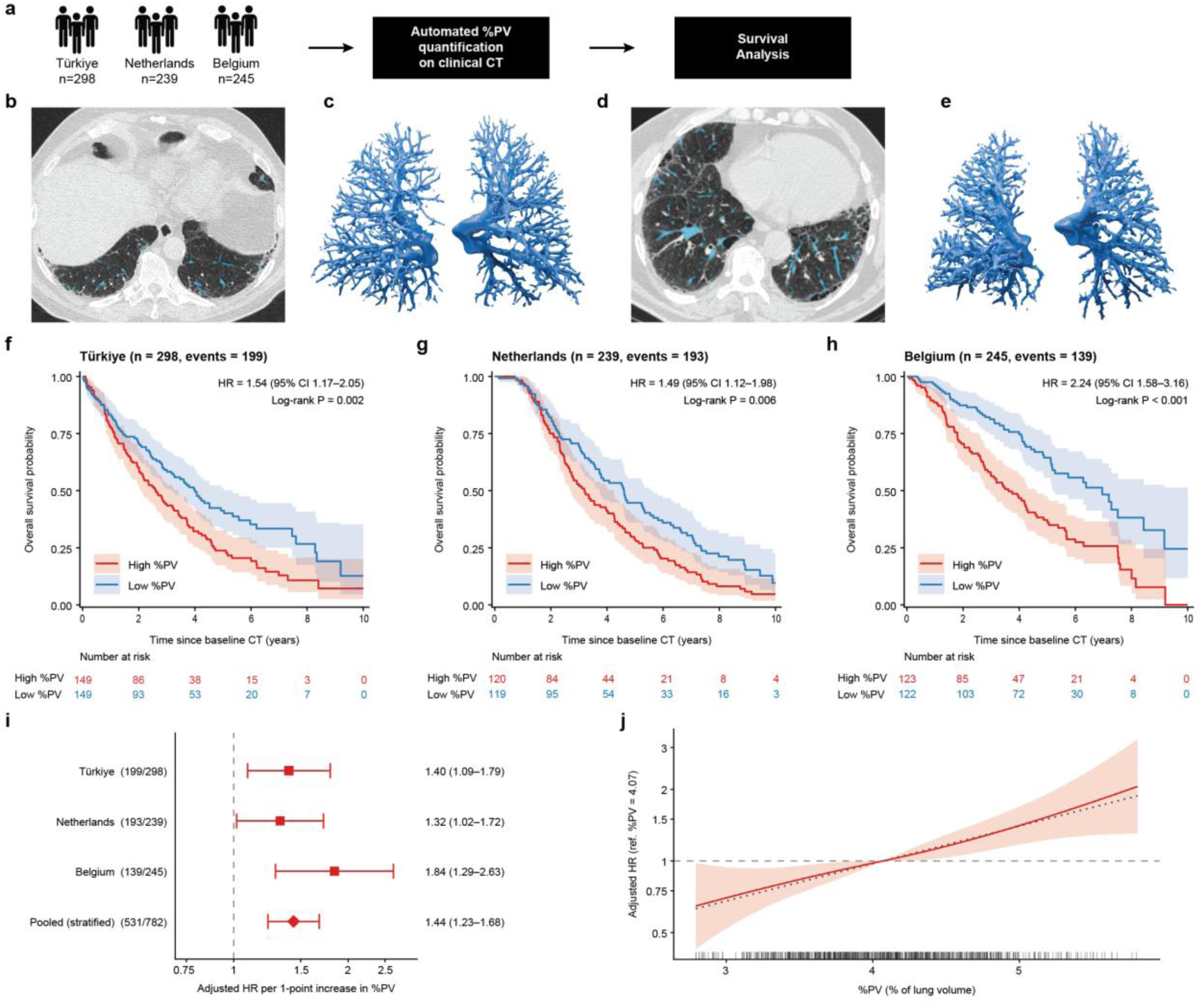
Quantification of intrapulmonary venous volume as a percentage of lung volume (%PV) on clinical CT scans and its association with survival. IPF patients from three tertiary centres (**a**) in Ege University Hospital, Izmir, Türkiye (n=298), St Antonius Hospital, Nieuwegein, Netherlands (n=239) and UZ Leuven, Belgium (n=245). Intrapulmonary venous volume was segmented on CT imaging using a radiologist-in-the-loop iterative Forced Perspective (nnU-Net-based) deep learning network with careful quality checking of model outputs. Axial CT sections with corresponding three-dimensional volumetric renderings of the pulmonary veins are shown (**b-e**) for two patients with IPF. The first axial CT (**b**) shows a 72-year-old male with limited IPF (baseline FVC 97% predicted) where 3.24% of the lung comprised pulmonary veins (**c**) who died 7.19 years after the CT. The second axial CT scan (**d**) shows a 68-year-old male with more extensive and severe IPF (baseline FVC 64% predicted) where 4.86% of the lung comprised pulmonary veins (**e**) who died 2.54 years after the CT. Kaplan-Meier survival curves show prognostic discrimination using the median value for the pulmonary vein percentage in the Turkish (**f**), Netherlands (**g**) and Belgian (**h**) clinical cohorts. Shaded areas indicate 95% confidence intervals, and numbers at risk are shown below each plot. Hazard ratios for the dichotomised %PV groups were obtained from unadjusted Cox proportional-hazards models fitted separately within each cohort, and P values from log-rank tests. (i) Adjusted hazard ratios for death per one-percentage-point increase in %PV in each cohort and in the pooled cohort. Cohort-specific models were adjusted for age, sex, smoking status, antifibrotic therapy, %FVC and %DLCO; antifibrotic therapy was omitted from the Belgian model because all patients received it. The pooled estimate (diamond) was obtained from the same multivariable model stratified by cohort. Numbers in brackets indicate deaths/total patients. Error bars indicate 95% confidence intervals, and symbol size reflects cohort size. %PV × cohort interaction P = 0.20; leave-one-cohort-out hazard ratios ranged from 1.37 to 1.54. (j) Adjusted dose-response relationship between %PV and all-cause mortality, modelled with a restricted cubic spline (3 degrees of freedom) in the primary multivariable model stratified by cohort and plotted between the 1st and 99th percentiles of %PV. The solid line shows the hazard ratio relative to the median %PV (4.07%), the shaded area the 95% confidence interval and the dotted line the linear model; tick marks on the x axis indicate individual patients. Test for non-linearity P = 0.85. In i and j, missing %FVC and %DLCO values were handled by multiple imputation (m = 20), and estimates were pooled using Rubin’s rules.

Multivariable Cox proportional-hazards modelling stratified by cohort (n=782, deaths=531) demonstrated that higher %PV was independently associated with all-cause mortality after adjustment for age, sex, smoking status, antifibrotic therapy, and disease severity as defined using forced vital capacity (%FVC) and diffusing capacity for carbon monoxide (%DLCO) (adjusted HR per one-percentage-point increase 1.44, 95% CI 1.23–1.68; P < 0.001; **Supplementary Table 3 and Supplementary Figure 13a**). The effect showed no relevant heterogeneity across cohorts (Fig. 7i), with no significant %PV × cohort interaction (P = 0.20), and was robust to sequential single-cohort exclusion (%PV hazard ratio 1.37 to 1.54 across the three leave-one-out analyses, all P < 0.001; **Supplementary Note 2**). The association was approximately linear across the observed range of %PV, with no evidence of a threshold (Fig. 7j), and was consistent across subgroups defined by age, sex, smoking status, GAP stage, %FVC and %DLCO (all interaction P > 0.1; Supplementary Figure 13b), including patients with GAP stage I disease (HR 1.43, 95% CI 1.10–1.85).

Consistent with the pooled analysis, cohort-specific Kaplan-Meier curves dichotomised at the within-cohort median of %PV demonstrated significantly shorter survival in patients with high %PV in all three cohorts (**Fig. 7f-h**; Türkiye HR 1.54, 95% CI 1.17–2.05; Netherlands HR 1.49, 95% CI 1.12–1.98; Belgium HR 2.24, 95% CI 1.58–3.16; all log-rank P ≤ 0.006). Quantitative CT measurement of intrapulmonary venous volume therefore identified venous expansion as an independent prognostic marker in IPF, distinct to underlying disease severity, extending the graded venous remodelling resolved at the organ scale by HiP-CT to a clinically measurable correlate of survival.

## DISCUSSION

Our study provides the first multiscale, multimodal demonstration that bronchial venous remodelling is an early and clinically relevant feature of IPF, linking endothelial reprogramming and spatial tissue organisation to three-dimensional vascular remodelling and patient outcome. Strikingly, bronchial venous endothelial cells extend beyond their normal peribronchial niche even in areas of mild fibrosis, indicating that this remodelling begins earlier in disease progression than previously appreciated. These cells acquire angiogenic, inflammatory, and extracellular matrix-remodelling phenotypes, establishing interactions with immune cells and fibroblasts at a specialised fibrovascular-immune interface. Together, these findings identify venous expansion as a previously overlooked component of fibrovascular remodelling in IPF, with implications for disease progression and prognosis.

Despite extensive clinical investigation, the involvement of the venous system in IPF pathobiology remains incompletely understood. In contrast to the pulmonary arterial circulation — where Liebow and colleagues described bronchial artery enlargement within fibrotic lungs as early as 1953^31^, and Turner-Warwick demonstrated precapillary anastomoses between the bronchial systemic and pulmonary arterial circulations in the 1960s^32^ — evaluation of the venous circulation has lagged. Murata and colleagues used microradiography and serial histological sections to characterise the anatomy of the bronchial venous plexus and its connections to the pulmonary veins in health^9^. Yet such studies of the pulmonary microvasculature have been limited by an inability to examine large three-dimensional volumes of lung tissue — a capability necessary in a spatially heterogeneous disease such as IPF, in order to understand how remodelling of the complex pulmonary and bronchial venous circulations develops across the spectrum of fibrotic severity.

Over the past decade, quantitative CT analysis has attempted global evaluation of the pulmonary vasculature, but cannot resolve individual vascular compartments or reliably distinguish vessels from fibrosis — a technique combining high spatial resolution with an extensive field of view is needed. HiP-CT delivers this step change in acquisition, but the resulting datasets are too large for manual annotation, and no established methodology existed to label vessels continuously across whole-organ and zoom imaging scales. We therefore developed a bespoke computer vision segmentation approach to visualise and quantify venous perturbations across regions of mild and severe fibrosis, directing our subsequent molecular analyses and enabling algorithms to quantify the pulmonary veins on clinical-scale imaging. More broadly, this work demonstrates that integrating whole-organ imaging with spatial and single-cell molecular profiling bridges biological scales to reveal mechanisms of chronic tissue remodelling invisible to any single modality alone.

Although the physiological role of venous remodelling remains to be determined, our data indicate that expansion of the bronchial venous circulation is accompanied by active endothelial reprogramming rather than passive vascular dilation. VWA1⁺/PLVAP⁺ bronchial vEC acquired angiogenic, inflammatory and matrix-remodelling programmes that persisted from mild to severe fibrosis and were independently reproduced by spatial transcriptomics. At the same time, capillary endothelial populations progressively declined and activated stress-response pathways consistent with the well-described capillary rarefaction of IPF. These findings suggest that endothelial remodelling reflects lineage-specific adaptation of distinct vascular compartments rather than a uniform endothelial response to fibrosis.

Beyond defining the endothelial lineage underlying venous remodelling, our data identify a specialised fibrovascular microenvironment centred on VWA1⁺/PLVAP⁺ bronchial vEC. These cells co-localised with fibroblasts, lymphatic endothelial cells and lymphoid cell populations, while spatial ligand–receptor analysis demonstrated a progressive transition from homeostatic endothelial–stromal communication towards matrix-rich, angiogenic and ultimately immune-regulatory signalling in advanced fibrosis. Mayr and colleagues recently described lymphoid aggregates associated with bronchial vessels in end-stage IPF^29^. Our findings refine this concept by demonstrating that this vascular-immune niche extends beyond organised tertiary lymphoid structures to accompany a fibrovascular interface associated with bronchial venous cells more broadly that is established already in mild fibrosis. These observations position bronchial venous EC as organisers of a spatially and molecularly integrated fibrovascular niche that coordinates extracellular matrix remodelling, vascular activation and immune cell activation and recruitment during disease progression.

However, several limitations should be acknowledged. The cross-sectional design precludes determining whether capillary rarefaction and bronchial venous expansion are mechanistically linked or represent parallel responses to shared fibrotic injury. Future functional studies will be needed to establish whether bronchial venous endothelial remodelling actively drives fibrosis or is an adaptive response to tissue remodelling. Likewise, we cannot distinguish whether expansion of the bronchial venous compartment reflects recruitment of pre-existing bronchial venous endothelium or acquisition of a bronchial venous programme by resident pulmonary endothelial cells, as previously observed ex vivo in human precision-cut lung slices after fibrotic cocktail stimulation^23^. The ex vivo and molecular analyses were performed in a small number of lungs; however, the organ-scale imaging phenotype was tested in 782 patients across three independent clinical cohorts, and the ex vivo findings were consistent across modalities with limited interpatient variation. Finally, neither HiP-CT nor clinical CT directly resolves endothelial lineage. Attribution of organ-scale venous remodelling to the systemic/bronchial compartment therefore relies on integration with the molecular datasets. Relatedly, the bronchial and pulmonary veins form an anatomically continuous drainage pathway that neither modality can separate. Clinical CT resolves only the pulmonary venous compartment — the bronchial veins fall below its detection limit, as they do on the downsampled whole-lung HiP-CT reconstructions used for segmentation — so %PV does not directly report the VWA1⁺/PLVAP⁺ bronchial venous endothelial phenotype. Nevertheless, the convergence of independent imaging (across multiple scales), spatial, and single-cell analyses provides strong evidence that bronchial venous endothelial remodelling is a central and clinically relevant feature of IPF progression.

In conclusion, by integrating non-destructive whole-organ microscopic-scale imaging, artificial intelligence, spatial and single-cell molecular profiling, and clinical imaging, we connect cellular and microanatomical disease processes directly to organ-scale pathology and patient outcome. In IPF, this approach uncovers venous remodelling as an early and clinically consequential feature of fibrosis, revealing an underappreciated vascular dimension of disease pathogenesis. More broadly, our study establishes a strategy for translating biological discoveries made in intact human organs into measurable phenotypes in living patients that could serve as a template approach with potential relevance well beyond pulmonary fibrosis.

## METHODS

### Ex vivo imaging: sample processing, scanning and reconstruction

Whole lungs were collected immediately following lung transplantation from patients diagnosed with IPF, fixed by infusion of 4% neutral buffered formalin via the main stem bronchus^18,23^. Control lungs were obtained as organ donations from patients undergoing euthanasia or whole-body donors from the Laboratoire d’Anatomie des Alpes Françaises in Grenoble, France. Lungs were collected with approval from the ethics committees of CHU UCL Namur, Mont-Godinne (B300202100073), Hôpital Erasme, Brussels (SBR2024109) and Hannover Medical School (10194_BO_K_2022). Prior to imaging, each specimen was prepared for synchrotron scanning according to a common protocol. Specimens were immersed in either 4% formalin or 70% ethanol as the surrounding medium, and the airways were filled with a gelled solution to provide internal structural support, using 0.2% Carbopol® 980 polymer gel for formalin-immersed specimens and a 70% ethanol gel for ethanol-immersed specimens. Each specimen was then immobilised within a container using layers of supporting material equilibrated with the surrounding medium. Residual air was removed from the tissue by degassing; during inline degassing, dissolved gas was continuously removed from the liquid surrounding the lung, promoting the progressive diffusion of trapped gas from the tissue into the surrounding medium. Both media were represented among the control and severe fibrosis specimens, so the surrounding medium was not confounded with fibrosis severity.

Specimens were scanned at the European Synchrotron Radiation Facility (ESRF) on beamline BM18, using a filtered white beam from a 1.1 T lateral-field source with a half-acquisition geometry. Whole-organ scans were acquired at isotropic voxel sizes of 16.6–42.3 µm, with average X-ray energies of 73–116 keV and propagation distances of 20–38 m; targeted zoom scans of regions of interest were acquired at 2.5–5.1 µm, with average X-ray energies of 81–117 keV and propagation distances of 1.5–6.0 m. Between 6,000 and 18,000 projections were recorded per scan. Tomographic reconstruction was performed with ring-artefact correction, followed by 16-bit conversion and JPEG-2000 compression. To make segmentation of the complete organ computationally tractable, whole-lung volumes were downsampled to 100 µm and zoom volumes to 17–20 µm prior to analysis; all reported measurements refer to these analysis volumes.

A separate set of lungs of patients with IPF leading to lung transplantation were air-inflated and fixed in the fumes of liquid nitrogen. These lungs were sliced with a bandsaw and systematically sampled using a power drill. Extracted samples were scanned in the frozen state on a FleXCT micro-CT scanner (imec-Visionlab, University of Antwerp) at a tube voltage of 45 kV and a target current of 442 µA, yielding isotropic voxels of 15 µm. These samples were divided into 2 and one piece was fixed in paraformaldehyde and embedded in paraffin for histology, spatial transcriptomics and multiplex immunofluorescence staining. The other half of the same sample was used for single nuclear sequencing. Control lungs were patients that underwent lobectomy for primary lung cancer. These patients had normal pulmonary function and no evidence of emphysema or fibrosis on CT. Tissue distant from the tumor was used. All lungs were collected with relevant ethical approval and signed informed consent in every hospital (UZA-EDGE001693). The stratification of mild vs severe fibrosis was based on micro-CT-based surface density measures performed with CTan. This was previously validated as a reliable readout of local fibrosis by demonstrating a strong association with Ashcroft scoring. The separation of mild vs severe fibrosis for the GeoMx analysis was performed based on local modified Ashcroft scoring by an expert thoracic pathologist where mild fibrosis was defined as an Ashcroft score ≤5 and severe fibrosis as a score >5^13,33^.

### Segmentation

We manually segmented pulmonary arteries and pulmonary veins from two whole lung HiP-CT scans resampled to 100µm isotropic voxel size using Thermo Scientific Amira Software which was used to develop a dedicated framework for expanding the incomplete manual segmentations and to label other HiP-CT data based on a Forced Perspective technique for three rounds^12^. Human observers (radiologists: D,Y., R.E., J.J.) reviewed the segmentations after each round of Forced Perspective before retraining models, with manual correction where applicable, in an active learning manner.

### Ex vivo venous volume quantification and statistical analysis

For each zoom volume of interest (VOI), analysed at a binned voxel size of 17-20µm, venous volume was quantified from the segmented venous trees as the combined volume of pulmonary and bronchial veins, expressed as a percentage of total sample volume. Although the bronchial (systemic) veins are resolved within HiP-CT zoom volumes, they are anatomically continuous with the pulmonary veins, and no consistent boundary exists at which the two can be divided. At whole-lung resolution, the bronchial veins cannot be separately resolved. Therefore, throughout the manuscript, venous volume refers to the combined intrapulmonary venous drainage compartment, and no inference is drawn regarding the relative contribution of either compartment. Venous volume was compared across the three fibrosis severity groups (control, mild and severe; n = 3 VOIs per group). Because unequal variances across severity groups were plausible and the small number of VOIs per group precluded formal assessment of normality, group differences were evaluated using Welch’s one-way ANOVA, followed by pairwise Welch’s two-sample t-tests with Bonferroni correction for multiple comparisons. Analyses were performed in R (version 4.5.0) using the rstatix package, with two-sided P < 0.05 considered statistically significant.

### Vascular Corrosion Casting and Scanning Electron Microscopy

Lungs for corrosion casting were obtained from Hannover Medical School (ethics approval 10194_BO_K_2022). At the time of tissue collection, the different vessels were cannulated using an olive-tipped cannula. The vasculature was flushed with saline at body temperature, followed by fixation with 2.5% glutaraldehyde solution (pH 7.4; Sigma-Aldrich, Munich, Germany). Subsequently, a prepolymerized PU4ii resin (VasQtec, Zurich, Switzerland), mixed with hardener (40% solvent) and blue dye, was injected as the casting medium. After complete resin polymerization, the lung tissue was macerated in 10% KOH (Fluka, Neu-Ulm, Germany) at 40°C for 2 to 3 days^34^. The resulting vascular casts were rinsed with water, frozen in distilled water, and freeze-dried. Thereafter, the casts were sputter-coated with gold in an argon atmosphere and examined using a Philips ESEM XL-30 scanning electron microscope (Philips, Eindhoven, The Netherlands) operated at 15 kV and 21 µA. Intussusceptive angiogenesis was identified by the presence of small transcapillary holes measuring 2–5 µm in diameter^35^.

### Single nuclei RNA-seq profiling and analysis

Smashed frozen lung tissue from different donor samples were pooled (2–4 samples per pool), transferred into a C-tube, and mechanically homogenized using a GentleMACS Dissociator (Miltenyi Biotec) in 1 mL of lysis buffer containing 10 mM Tris-HCl (pH 7.5), 146 mM NaCl, 21 mM MgCl₂, 1 mM CaCl₂, 0.01% BSA, 0.03% Tween-20, 1 mM DTT, 10% EZ buffer (Sigma), 1 U/µL RNase inhibitor, and nuclease-free water. To terminate lysis, 2 mL of wash buffer (10 mM Tris-HCl, pH 7.5; 10 mM NaCl; 3 mM MgCl₂; 1% BSA; 1 mM DTT; 0.5 U/µL RNase inhibitor) was added. Samples were centrifuged at 300 × g for 5 min at 4°C, the supernatant removed, and the pellet resuspended in 900 µL wash buffer. The suspension was filtered sequentially through a 5 µm Mini strainer (PluriSelect), centrifuged under the same conditions, and the pellet resuspended in 900 µL wash buffer for a second filtration through a 5 µm strainer. After a final centrifugation, the nuclei pellet was resuspended in 100 µL of resuspension buffer (10 mM Tris-HCl, pH 7.5; 10 mM NaCl; 3 mM MgCl₂; 1% BSA; 1 mM DTT; 1 U/µL RNase inhibitor) and counted. Isolated nuclei were then processed for single-cell encapsulation targeting 10 000 cells, followed by library preparation and sequencing on the Illumina NovaSeq 6000 platform, in accordance with the manufacturer’s protocol (10x Genomics, CG000315 RevE). Genomic DNA was isolated from 13 donors and genotyped using the Illumina infinium global screening array.

Mapping and counting of the UMIs for the samples was performed using 10x Genomics Cell Ranger (version 9.0.0) with the reference genome GRCh38-2020-A provided by 10x Genomics^36^. Souporcell (version 2.5.0) was used for donor deconvolution of the pools. The resulting cluster genotypes called by Souporcell were then compared to the SNP array data to match the deconvoluted snRNA-seq clusters back to the original donor^37^. Three different tools were used for this: bcftools stats (version 1.12), plink2 --sample-diff (version 2.0), and SoupLadle (version 0.0.0.9000)^38–40^. All deconvoluted clusters could be matched back to a single donor and all three tools were in agreement. snRNA-seq analyses up to differential gene expression analysis were performed in R version 4.4.1. Ambient RNA contamination was removed using the R package SoupX (version 1.6.2) with a fixed contamination fraction of 0.3^41^. Doublets were detected with the package scDblFinder (version 1.20.0) using the cluster-based approach^42^. The Scater package (version 1.34.0) was used to assess the proportion of ribosomal and mitochondrial genes as well as the number of detected genes^43^. Cells were considered as outliers and filtered out if the value of the proportion of expressed mitochondrial genes or the number of detected genes deviated more than three median absolute deviations from the median across all cells. Additionally, cells where Souporcell could not unambiguously assign a single cluster were filtered out. Normalization between samples was done with the sctransform method of Hafemeister and Satija using the package Seurat (version 5.1.0)^44,45^.

After filtering out doublets identified by scDblFinder and other low-quality cells, the samples were batch corrected in Python (version 3.12.7) with the scVI model of the scvi-tools library (version 1.3.0)^46,47^. The model was trained using a latent space with 30 dimensions, 2 hidden layers and the dispersion setting “gene-batch” for a maximum of 70 epochs with the possibility of early stopping. Other parameters were left at default values.

### DGE and pathway analysis

Differential expressed genes between conditions (overall and within each clusters) were identified with the FindMarkers function of Seurat in R^45^. The MAST package (version 1.32.0) was used for the differential testing with donor as a latent variable. Genes were considered upregulated if they met the criteria of *P* value < 0.05 and log2 fold change (log2FC) > 0.3, while downregulated genes were defined as those with *P* value < 0.05 and log2FC < −0.3^48^. Functional enrichment analysis of upregulated and downregulated genes was performed using over- representation analysis (ORA) with the Gene Ontology Biological Process 2025 database (GO_Biological_Process_2025) implemented in GSEApy (version 1.1.10). Pathways with an enrichment significance cutoff of *P* < 0.05 were considered significantly enriched^49^. Upregulated and downregulated biological processes were identified separately based on the directionality of gene expression changes.

### Augur-based pertubation analysis

Augur-based analysis was conducted using Pertpy (version 1.0.1) to prioritize cell types according to their perturbation responsiveness^25,50^. A Random Forest classifier was applied at the Level_2 meta cell type annotation to compare IPF mild versus control and IPF severe versus control conditions. The resulting mean Augur scores were aggregated into a matrix for downstream interpretation.

### Cell–cell communication analysis with Cell Chat analysis

Cell–cell communication analysis was performed using CellChat (version 2.2.0) to compare signaling interactions across conditions^30^. The analysis incorporated the complete human CellChat database, excluding non-protein signaling interactions. Ligand–receptor interactions were inferred using the default statistical framework implemented in CellChat. Mesothelial cells were removed from the analysis due to their low abundance. To improve signal detection and reduce noise from weak interactions, analyses were performed at the meta cell-type level by aggregating related cell populations. VWA1⁺PLVAP⁺ bronchial vEC were defined as the source population to investigate outgoing communication patterns. Differential signaling between conditions was assessed by comparing communication probabilities for each ligand–receptor pair, with increases or decreases defined based on relative changes in inferred interaction strength across groups.

### Iterative indirect immunofluorescence imaging (4i) and analysis

Tissue microarray (TMA) formalin-fixed paraffin-embedded (FFPE) sections of IPF and control lung samples were baked at 60 °C for 1 hour. Sections were then deparaffinized in xylene (2 × 10 min) and rehydrated through a graded ethanol series (2 × 100%, 90%, 80%, and 70%) before rinsing in distilled water for 2–5 min. Antigen retrieval was performed using R-Universal Buffer (Aptum; diluted 1:10 in Milli-Q water) in a pressure cooker for 20 min. Slides were allowed to cool to room temperature and subsequently washed twice in PBS. To reduce nonspecific binding, sections were incubated with 150 mM maleimide (Merck) prepared in 1% BSA for 1 hour at room temperature. Sections were then incubated overnight at 4 °C with primary antibodies (Supplementary Table 5) diluted in PBS containing 1% BSA^22^. The following day, slides were washed and incubated with appropriate secondary antibodies (Invitrogen or Lubio) and DAPI (1:250 dilution) in PBS supplemented with 1% BSA for 2 hours at room temperature (Supplementary Table 5). Autofluorescence was quenched using the Vector Autofluorescence Quenching Kit (Vector Laboratories; reagents A:B:C mixed at 1:1:1) for 2 min in the dark. An imaging buffer consisting of 700 mM N-acetylcysteine (Merck), 20% HEPES, and 80% ddH₂O was applied to the sections prior to imaging. Whole-slide imaging was performed using a Hamamatsu NanoZoomer S60 slide scanner. Following imaging, antibodies were stripped by incubating the sections in elution buffer containing 0.5 M L-glycine (Merck), 3 M urea (Merck), 3 M guanidine hydrochloride (Merck), and 70 mM TCEP-HCl (Merck) prepared in 20 mL ddH₂O. Elution was performed three times for 20 min each. After elution, sections were incubated with imaging buffer and examined under a microscope to confirm complete removal of fluorescence signal. Upon verification, subsequent staining cycles were initiated by repeating the blocking and antibody incubation steps with new antibody panels.

Images acquired from multiple staining cycles were converted to OME-TIFF format for downstream analysis. Background signal was reduced using the open-source software PyImageJ (version 1.8.0)^51^, followed by image alignment across iterative staining cycles using open-source software image_registration_tool (version 0.1.0) to ensure spatial correspondence. Cell segmentation was performed in QuPath (version 0.6.0) using the watershed cell detection algorithm with the following parameters: background radius = 20.0 µm, background by reconstruction = true, median radius = 0.0 µm, sigma = 0.6 µm, minimum area = 15.0 µm², maximum area = 90.0 µm², threshold = 15.0, watershed post-processing = true, cell expansion = 2.0 µm, nuclei included = true, boundary smoothing enabled, and measurement calculation disabled^52^. Quality control was conducted using SPACEc (version 0.1.2)^27^, where approximately 1% of cells were filtered out based on abnormal DAPI intensity and cell size to remove potential artifacts and segmentation errors. Following quality filtering, the data were normalized using a z-score normalization approach. The normalized dataset was subsequently analyzed using Squidpy (version 1.6.5) for downstream spatial and single-cell analyses^53^.

Cellular neighborhood analysis was performed using SPACEc (version 0.1.2), where cells were grouped into eight distinct neighborhoods based on their spatial organization and phenotypic profiles. Patch proximity analysis was also conducted using SPACEc (version 0.1.2) to assess spatial relationships between cellular clusters. A minimum cluster size of 5 cells was applied to define patches for proximity analysis.

### Spatial transcriptomics

The Bruker GeoMx DSP platform (Seattle, USA) was used in conjunction with the whole transcriptome atlas (WTA)^21^. FFPE specimens were immobilized onto slides by drying at 60°C for 3 hours prior to the assay. Samples were deparaffinized through a descending solvent series: three 5-minute washes in xylene, followed by two 5-minute washes in 100% ethanol, one 5-minute wash in 95% ethanol, and a 1-minute rinse in 1× PBS. The slides were then incubated in 1× Tris-EDTA (pH 9.0) in a pressure cooker at low pressure for 20 minutes at 99°C for antigen retrieval. Proteinase K treatment in PBS was applied at 37°C for 15 minutes to further retrieve the targets. After washing thoroughly with PBS, samples were postfixed in 10% neutral buffered formalin (NBF) for 5 minutes, followed by two 5-minute washes in NBF stop buffer and a 5-minute wash in PBS. Next, GeoMx RNA probes were hybridized overnight (16-17 hours) at 37°C in a ZytoBrite Hybridizer. PBS was replaced with 200 μL of hybridization solution containing the probe mix (WTA Whole Transcriptome Atlas Human RNA, Bruker) per slide, which was then covered with a HybriSlip (Grace Biolab). After incubation, unbound probes were removed by washing in a pre-warmed stringent wash solution containing 50% formamide and 2× SSC. Coverslips were removed by submerging the slides in washing solution, followed by two 25-minute washes at 37°C and two 2-minute washes in 2× SSC at room temperature. Slides were blocked with buffer W for 30 minutes in a humidified, light-protected chamber. The blocking buffer was replaced with 200 μL of an antibody mix, which was incubated for 1 hour at room temperature in the dark. Anti-Pan cytokeratin Alexa488 (Novus Biologicals, NBP2-33200, 1:500), anti-CD31 Alexa594 (Proteintech, CL594-11265, 1:40) and anti-CD90 (Proteintech, 66766-1-Ig, 1:40) antibodies were used, along with SYTO83 (Invitrogen, S11364, 1:25,000). Anti-CD90 was labelled with Alexa647 using the labelling kit (Thermo Fisher Scientific, A20186) according to the manufacturer’s instruction. After incubation, the slides were washed and stored in 2× SSC. The slides were then washed twice for 5 minutes in 2× SSC before being placed in the GeoMx DSP slide holder with 3 mL of buffer S. The slides were scanned on the GeoMx DSP platform, with constant exposures of 100 ms for Cytokeratin, CD31 and CD90, and 50 ms for SYTO83. Regions of interest (ROIs) were defined, and oligonucleotides were photo-released. The DSP samples were then collected in a 96-well PCR plate for downstream processing. The 96-well plates were sealed with air-permeable foil and allowed to dry completely overnight under a hood. Next, the DSP samples were rehydrated with 10 μL of DEPC-treated water, covered with an adhesive seal, and incubated at room temperature for 10-20 minutes. Library preparation was performed in a new 96-well PCR plate, using 2 μL of GeoMx NGS Master Mix, 4 μL of Seq Code Primer Mix (from the corresponding 96-well plate), and 4 μL of DSP sample from the collection plate using a MiniAmp Thermal Cycler (Applied Biosystems) with a 100°C heated lid. After PCR amplification, 4 µL of all samples, including internal no-template controls (NTC), were collected into a single 1.5 mL tube and purified using magnetic beads under nuclease-free conditions. The library quality was assessed via TapeStation analysis, with the characteristic peak at 170 bp. Next-generation sequencing (NGS) was performed using the NovaSeq 6000 platform, with 250 pM of library and 5% spike-in as an internal control. FASTQ files were initially processed using the GeoMx NGS Pipeline (Bruker) to convert raw sequencing data into a usable format for analysis. Following this, quality control (QC) measures were applied to ensure data integrity and reliability. The primary QC thresholds included a minimum of 1,000 raw reads per region of interest (ROI), a sequencing saturation rate greater than 50%, and a minimum of 80% of the reads being aligned, stitched, and trimmed. Additionally, ROIs that exhibited expression of fewer than 10% of the genes were excluded from further analysis to ensure the inclusion of only biologically meaningful data. No segments failed QC metrics. To further refine the dataset, genes that were undetected in 10% or more of the ROIs were removed. This process resulted in a gene list of 18,677 genes. Finally, raw count data underwent negative RNA probe-based quantile normalization (Q3 normalization).

### Endothelial Cell Deconvolution

Endothelial cell subtype composition within GeoMx Digital Spatial Profiling (DSP) CD31^+^ regions was estimated using reference-based transcriptomic deconvolution with CIBERSORTx (https://cibersortx.stanford.edu/)^19,20^. A custom endothelial cell signature matrix was generated based on the Lung Cell Endothelial Atlas published by Schupp et al.^15^, which defines transcriptional profiles of distinct pulmonary endothelial cell populations. The reference signature matrix was constructed from single-cell RNA sequencing data by selecting endothelial cell clusters and representative marker genes that capture cell-type-specific expression patterns. Normalized gene expression data from GeoMx CD31^+^ regions were used as the mixture matrix for deconvolution. CIBERSORTx was run using the provided endothelial signature matrix with the relative mode to estimate the relative abundance of endothelial cell subpopulations within each spatial region. Analyses were performed using the custom signature matrix framework implemented in CIBERSORTx, with quantile normalization disabled due to the use of GeoMx DSP expression data and batch correction applied as recommended for cross-platform deconvolution analyses. Statistical significance of deconvolution estimates was assessed using CIBERSORTx permutation testing with 1,000 permutations. Output fractions were normalized to sum to one within each CD31^+^ region and used for downstream comparative analyses of endothelial cell subtype representation across experimental groups.

### Ligand-receptor interaction analysis between CD31^+^ and CD90^+^ GeoMx regions

Potential cell-cell communication between endothelial-enriched CD31^+^ regions and adjacent CD90^+^ stromal regions was inferred using LIANA (Ligand-Receptor ANalysis framework) implemented in Python^28^. Normalized gene expression matrices from GeoMx regions were converted into an AnnData object and annotated according to spatial compartment identity, with CD31^+^ regions representing endothelial-enriched areas and CD90^+^ regions representing adjacent stromal compartments. Because GeoMx DSP regions represent multicellular spatial units rather than individual cells, each region was treated as a spatially resolved communication unit for ligand–receptor inference. LIANA was applied to identify and rank potential ligand–receptor interactions between CD31^+^ and CD90^+^ compartments using integrated ligand–receptor resources, including the CellPhoneDB and ConnectomeDB-derived interaction databases^28,54,55^. The analysis incorporated ligand and receptor expression levels across spatial compartments and generated consensus interaction scores based on multiple ligand–receptor inference methods. Significant interactions were identified based on LIANA ranking metrics and statistical significance criteria, with interactions retained if they were supported by expressed ligand–receptor pairs and passed a predefined significance threshold (adjusted P value <0.05). Directional communication analyses were performed separately for endothelial-to-stromal (CD31^+^ to CD90^+^) and stromal-to-endothelial (CD90^+^ to CD31^+^) signaling.

### Pathway enrichment analysis and network visualization using Metascape

Functional enrichment analysis was performed using Metascape (https://metascape.org/) to identify biological pathways and molecular processes associated with differentially expressed genes identified within systemic venous endothelial cells between mild and severe regions^56^. Gene lists were uploaded to Metascape with species-specific annotation and pathway enrichment was performed using the integrated databases, including Gene Ontology (GO), Kyoto Encyclopedia of Genes and Genomes (KEGG) and Reactome. Significantly enriched pathways were identified based on statistical enrichment analysis, with pathways meeting predefined significance thresholds (adjusted P value <0.05) considered for downstream visualization. Enriched pathways were visualized as an enrichment network in which individual nodes represented significantly enriched biological terms and edges represented similarity relationships between pathways based on shared gene membership. Closely related terms were clustered into functional modules, and representative pathway terms were selected for visualization. To compare pathway contributions between severe and mild IPF groups, the proportion of gene enrichment attributable to each group was calculated for individual pathway nodes and displayed as a surrounding pie chart around each network node. The relative contribution of each disease severity group was represented by the fraction of genes associated with the corresponding pathway that originated from mild or severe regions. Network plots were generated using Metascape visualization outputs and further customized for graphical presentation.

### Clinical CT Analysis

#### Study Design and Patient Cohorts

This multicenter retrospective study included three independent cohorts of patients diagnosed with idiopathic pulmonary fibrosis (IPF) by multidisciplinary teams: Türkiye (n=298), the Netherlands (n=239) and Belgium (n=245). The final pooled analysis comprised 782 patients with 531 deaths. This study was approved by the institutional review boards by each participating centre and by the Leeds East Research Ethics Committee: 20/YH/0120. The requirement for informed consent was waived due to the retrospective nature of the study. All procedures were conducted in accordance with the Declaration of Helsinki and applicable local regulations. Lung function results including FVC and DLCO, obtained within 3 months of the CT scan were collected for use as covariates in survival models. Clinical data and follow-up information were also recorded. Patient selection across all three cohorts is summarised in Supplementary Figure 12.

#### CT acquisition and venous segmentation

All patients underwent baseline, volumetric, end-inspiratory, non-contrast chest CT imaging with a maximal slice thickness of 1mm. Quantitative analysis of pulmonary venous volumes was performed using a dedicated in-house developed variant of the Forced Perspective segmentation method specific to clinical CT scans^12^. Manual labels for training were produced by subspecialist thoracic radiologists (D.Y., R.E., J.J.). The segmentation framework itself was developed on HiP-CT data and adapted here to the resolution and contrast characteristics of clinical CT. Intrapulmonary venous volume (%PV) was calculated as a percentage of total lung volume.

#### Survival Analysis

The primary outcome was all-cause mortality. Lung transplantation was treated as a censoring event. Follow-up time was calculated from the date of baseline CT scan to the date of death, lung transplantation, or last follow-up. Patients who were alive and had not undergone lung transplantation at the end of follow-up were censored at the date of last contact. Cohort-specific transplant censoring rates are reported in Supplementary Note 1. The association of %PV and each covariate with mortality was first assessed in univariable Cox proportional-hazards models stratified by cohort (Supplementary Table 3). Hazard ratios (HR) with 95% confidence intervals (CI) were calculated per one-percentage-point increase in %PV. For display purposes, Kaplan– Meier survival curves were constructed separately for each cohort after dichotomising patients into high- and low-%PV groups at the cohort-specific median. Survival was compared between groups using the log-rank test, and unadjusted hazard ratios for the dichotomised groups were estimated from Cox proportional-hazards models fitted within each cohort. Number-at-risk tables are shown beneath each curve.

Multivariable Cox proportional hazards regression models stratified by cohort were constructed to evaluate the independent prognostic value of %PV after adjusting for established clinical predictors. The following covariates were included in all models: age at CT scan, sex, smoking status (never vs. ever), antifibrotic use, forced vital capacity (percent predicted = %FVC), and diffusion capacity for carbon monoxide (percent predicted = %DLCO). Missing values for smoking status, %FVC, and %DLCO were imputed using multiple imputation by chained equations (MICE) with twenty imputations^57^. Details of the imputation procedure, including the proportion of imputed values per cohort and the handling of a single missing smoking-history record, are provided in Supplementary Note 1. The assumption of a linear relationship between %PV and the log hazard was checked before modelling by comparing the linear model with one in which %PV was fitted as a restricted cubic spline; there was no evidence of departure from linearity (likelihood ratio test, P = 0.85), so %PV was entered as a continuous linear term. The proportional hazards assumption of the multivariable stratified Cox model was assessed using scaled Schoenfeld residuals (cox.zph in the survival R package). For covariates that violated the assumption, a pre-specified remediation analysis was performed in which time-varying coefficients were added using a log-time transformation (Supplementary Note 3).

Subgroup analyses were performed to assess the consistency of the association between %PV and mortality across clinically relevant strata: age (<70 vs. ≥70 years), sex, smoking status, GAP stage (I, II and III, calculated from age, sex, %FVC and %DLCO), %FVC (≥80% vs. <80% predicted) and %DLCO (split at the pooled median, 46.6% predicted). Within each subgroup, the multivariable model stratified by cohort was fitted with the same covariates, omitting the defining variable for sex and smoking status. Heterogeneity was assessed by adding a %PV × subgroup interaction term to the model fitted to the full cohort; for GAP stage, the two interaction terms were tested jointly using a multivariate Wald test pooled across imputations. Subgroups defined by imputed variables were assigned within each imputed dataset, and estimates were combined using Rubin’s rules.

#### Cohort consistency and sensitivity analyses

Consistency of the %PV effect across cohorts was evaluated within the patient-level stratified Cox framework. A likelihood-ratio test compared the primary model with an extended model that additionally included PV × cohort interaction terms; a non-significant result was interpreted as supporting a common %PV effect across cohorts. As a within-cohort sensitivity description, cohort-specific multivariable Cox models were additionally fitted (Supplementary Table 4). Cross-cohort robustness was assessed by patient-level leave-one-out analysis: the multivariable stratified Cox model was re-fitted after sequentially excluding each cohort, and the %PV hazard ratio compared across iterations (Supplementary Note 2).

### Usage of Generative AI

Generative AI tools (ChatGPT, OpenAI; Claude, Anthropic) were used to assist with the development, debugging and refinement of code for data analysis and visualization. All AI-assisted code was reviewed, tested and validated by the authors, and all analyses and scientific interpretations were performed and verified by the authors.

### Statistical Analysis

All statistical analyses for the clinical CT cohorts were performed using R version 4.6.1 (R Foundation for Statistical Computing, Vienna, Austria). The survival package was used for Cox regression and Kaplan-Meier analyses, and figures were produced with ggplot2 (version 4.0). Two-sided P values less than 0.05 were considered statistically significant. For the omics data, comparisons among the three groups were performed using the two-sided Kruskal–Wallis test, followed by Dunn’s post hoc test with Bonferroni correction for pairwise comparisons.

## Data Availability

Raw and processed GeoMx DSP data has been uploaded to the GEO database (GSE316287).

The raw snRNA-seq data have been deposited in the Gene Expression Omnibus (GEO) under accession number GSE347725. The annotated snRNA and 4i datasets are available on Zenodo: https://doi.org/10.5281/zenodo.22685651

The HiP-CT datasets were acquired under ESRF beamtimes md1290 and md1389 and will be made publicly available through the Human Organ Atlas (https://human-organ-atlas.esrf.fr) upon publication; dataset DOIs will be provided in the final version of the manuscript.

The clinical CT imaging and associated clinical data from the three patient cohorts cannot be made publicly available because the consent obtained and the applicable national regulations do not permit open sharing of patient imaging data. De-identified data may be made available for academic research from the corresponding author (J.J.) upon reasonable request, subject to approval from the contributing centres and the relevant ethics committees and to a data use agreement.

## Code Availability

The algorithms and custom code devised for image analysis during the current study are not publicly available due to ongoing commercialization and patent-pending status. The code may be made available from the corresponding author upon reasonable request and subject to execution of a material transfer agreement and/or non-disclosure agreement. The scripts used for snRNA-seq and 4i spatial analysis are available on GitHub:

https://github.com/gote-schniering-lab/2026_Yamada_Wang_et_al_Multiscale_mapping_of_venous_remodelling_in_IPF.git

## Funding

This publication has been made possible in part by CZI grant 2022-316777 and CZIF2024-009938 from the Chan Zuckerberg Initiative DAF, an advised fund of Silicon Valley Community Foundation (funder DOI 10.13039/100014989), in part by grant number CZIF2021-006424 from the Chan Zuckerberg Initiative Foundation, the Rosetrees Trust (PGL25-Full/100046, CF-2023-M-2\105) and the Robert Luff Foundation (PGL25-Full/100046), the NIHR UCLH Biomedical Research Centre and in whole or in part by the Wellcome Trust ( 209553/Z/17/Z and 227835/Z/23/Z). This project was partially funded by German Center for Lung Research (DZL). We gratefully acknowledge ESRF beamtimes md1290 and md1389 on BM18 as sources of the data. DY is supported by grants from the fellowship of Astellas Foundation for Research on Metabolic Disorders. SA was supported by the International Alliance for Cancer Early Detection, an alliance between Cancer Research UK [EDDAPA-2023/100002], Canary Center at Stanford University, the University of Cambridge, OHSU Knight Cancer Institute, University College London and the University of Manchester. JB received funding from the Chan Zuckerberg Initiative Foundation under Grants CZIF2021-006424 and CZIF2022-316777, the UK Medical Research Council (MR/R025673/1), and the Royal Academy of Engineering (Ciet1819-10). JGS received funding from the Swiss National Science Foundation (grant number: 234552), Swisslife and Lungenliga Bern foundation. The funders had no role in study design, data collection and analysis, decision to publish or preparation of the manuscript.

## Competing interests

R.E. has received lecture fees from Nippon Boehringer Ingelheim. M.Ver. received institutional support from Boehringer Ingelheim and support for meeting attendance from Boehringer Ingelheim and Sanofi. CW has received speaker fees from Boehringer Ingelheim. SEV received funding from Chiesi NV not related to this work and grant funding via the special research fund of the University of Antwerp and the Collen Francqui foundation. J.J. declares consultancy fees from Boehringer Ingelheim, F. Hoffmann-La Roche, Open Source Imaging Consortium, GlaxoSmithKline, Voiant, NHSX; fees from advisory Boards for Boehringer Ingelheim, F. Hoffmann-La Roche, GlaxoSmithKline; lecture fees from Boehringer Ingelheim, F. Hoffmann-La Roche, Takeda; grant funding from GlaxoSmithKline, Wellcome Trust (209553/Z/17/Z and 227835/Z/23/Z), EU Horizon 2020, Microsoft Research, Gilead Sciences, Chan Zuckerberg Initiative (CZIF2024-009938), Rosetrees Trust (PGL25-Full/100046, CF-2023-M-2\105) and the Robert Luff Foundation (PGL25-Full/100046). BM declares research grants from AbbVie, Protagen, and Novartis Biomedical Research; declares a mir-29 patent for the treatment of systemic sclerosis (US8247389, EP2331143); declares lecturing fees from Boehringer Ingelheim, GlaxoSmithKline, Novartis, Otsuka, Merck Sharp & Dohme, and Lilly; declares consulting fees from Novartis, Boehringer Ingelheim, Janssen-Cilag, and GlaxoSmithKline; declares financial support for congresses (e.g., registration fees and travel) from Medtalk, Pfizer, Roche, Actelion, Mepha, Boehringer Ingelheim, and Merck Sharp & Dohme; and has served on advisory boards for Janssen and Boehringer Ingelheim. All other authors declare no competing interests.

## Materials & Correspondence

Correspondence and requests for materials should be addressed to M.A., S.E.V., J.G.S. or J.J.

## Acknowledgments

Microscopy was performed on equipment supported by the Microscopy Imaging Center (MIC), University of Bern, Switzerland. We thank the Next Generation Sequencing Platform of the University of Bern for performing the high-throughput sequencing experiments. The Interfaculty Bioinformatics Unit (IBU), University of Bern provided computational infrastructure and support with bioinformatic analyses. Illumina SNP phenotyping experiments were performed at the iGE3 Genomics Platform of the University of Geneva (https://ige3.genomics.unige.ch).

We acknowledge the European Synchrotron Radiation Facility (ESRF) for the provision of synchrotron beamtime and thank the staff of beamline BM18 for their assistance (proposals md1290 and md1389).

We are grateful to Professor Kiyoshi Murata and Professor Harumi Itoh, whose earlier anatomical studies of the bronchial venous plexus informed this work, for reviewing the HiP-CT reconstructions with us and for their advice on the interpretation of the bronchial venous anatomy.

The authors sincerely thank those who donated their bodies to science so that anatomical research could be performed. Results from such research can potentially increase humanity’s overall knowledge that can then improve patient care. Therefore, these donors and their families deserve our highest gratitude. We thank the Laboratoire d’Anatomie des Alpes Françaises (LADAF), Université Grenoble Alpes, for providing access to the donor specimens.

## Author Contributions

D.Y., H.W., A.S., and J.R. contributed equally to this work. D.Y., R.E. and A.S. performed HiP-CT and clinical imaging, vascular segmentation and analysis. H.W. performed and analysed the single-nucleus RNA-sequencing and 4i experiments and led the bioinformatic analysis. J.R. performed and analysed the GeoMx spatial transcriptomic experiments and contributed to the integrated spatial analyses. The four co-first authors contributed to data interpretation, figure preparation and preparation of the manuscript. J.J., J.G.S., S.E.V. and M.A. contributed equally to the supervision of this work. They conceived and designed the study, supervised the experimental and analytical work, integrated and interpreted the findings across modalities, and wrote and revised the manuscript. R.Bo. and M.B. contributed to single-nucleus RNA-sequencing and spatial proteomic experiments and analyses. J.B., T.U., J.P., H.D., P.T., C.L.W., P.D.L., S.A., K.E., and B.T. contributed to HiP-CT imaging, image processing and quantitative analyses. S.K.R. and A.F. contributed to histological validation of the vascular segmentation. D.C., Y.C., C.H.M.v.M., F.T.v.B., M.Vel., H.W.v.E., N.M., R.S., B.G., T.S.L., W.A.W., T.G., T.F., K.A., M.Ver., and L.J.d.S. contributed to clinical imaging analyses and clinical data acquisition. D.J., L.N., C.W., H.F., J.M.H.H., M.V.K., J.C.S., M.G.J., F.M.C., J.M.B., B.M.V., and S.D. contributed tissue samples, clinical data and associated pathological characterization. R.Br., S.K., M.K., B.M. contributed methodological, computational and/or analytical expertise and participated in data interpretation. All authors reviewed the manuscript and accepted the final version.

## DATA SUPPLEMENT

**Supplementary Movie 1. Whole-organ HiP-CT imaging of the venous drainage compartment in control and IPF lungs.**

Volumetric renderings of an intact control lung (left) and an explanted IPF lung (right), acquired by hierarchical phase-contrast tomography (HiP-CT) at isotropic voxel sizes of 16.6–42.3 µm and resampled to 100 µm for segmentation. The segmented venous compartment is shown in blue; surrounding parenchyma and airways are rendered in shades of copper-brown, with denser structures such as airway walls and pleura appearing brighter. In the control lung, venous branching follows a regular, hierarchically ordered course extending uniformly toward the periphery. In the IPF lung, the venous compartment is redistributed toward the subpleural and interlobular septal regions, with irregular dilatation of proximal segments and loss of the ordered peripheral branching seen in the control; honeycomb remodelling is evident at the lung base.

**Supplementary Note 1. Multiple imputation, smoking history handling, and lung transplant censoring**

Multiple imputation. Baseline pulmonary function values were considered paired with the baseline CT only when measured within three months of the scan. Missing values of forced vital capacity (FVC) and diffusing capacity for carbon monoxide (DLCO) were imputed using multiple imputation by chained equations (MICE) as implemented in the mice R package, with twenty completed datasets (m = 20). The imputation model included all variables used in the survival analysis (%PV, age, sex, smoking history, antifibrotic therapy exposure, FVC, DLCO, follow-up time, and the event indicator) so that imputed values remained consistent with the analytic relationships of interest. Missing FVC values were imputed for 52 of 298 (17.4%) in the Turkish cohort, 50 of 239 (20.9%) in the Netherlands cohort, and 29 of 245 patients (11.8%) in the Belgian cohort; missing DLCO values were imputed for 90 of 298 (30.2%), 51 of 239 (21.3%), and 37 of 245 (15.1%), respectively. The Turkish cohort had the highest proportion of imputed DLCO values, reflecting the higher prevalence of incomplete diffusion testing within the 93-day pairing window at that centre.

Smoking history. Smoking history was missing for a single Turkish patient (1 of 298; 0.3%) and was assigned the cohort-specific mode (ever-smoker, consistent with the 68.8% prevalence of ever-smokers in the Turkish cohort). A sensitivity analysis that excluded this patient (n = 781) yielded a multivariable %PV hazard ratio identical to two decimal places (HR 1.44, 95% CI 1.23–1.68; P < 0.001), indicating that the assignment did not influence the principal result.

Lung transplant censoring. Patients who underwent lung transplantation during follow-up were censored at the date of transplantation, regardless of their subsequent survival status, and therefore contributed to the at-risk population up to, but not beyond, that date. Transplantation occurred in 19 of 245 patients (7.8%) in the Belgian cohort, 23 of 239 (9.6%) in the Netherlands cohort and none in the Turkish cohort.

**Supplementary Note 2. Cross-cohort consistency and patient-level cross-validation**

The %PV effect was consistent across the three cohorts. In cohort-specific multivariable Cox proportional-hazards regression fitted independently in each cohort with adjustment for the same clinical covariates as the primary model, with antifibrotic therapy omitted from the Belgian model owing to 100% antifibrotic exposure — the cohort-specific hazard ratios for %PV (per one-percentage-point increase) were: Türkiye HR 1.40 (95% CI 1.09–1.79; P = 0.009; n = 298, 199 deaths); the Netherlands HR 1.32 (95% CI 1.02–1.72; P = 0.037; n = 239, 193 deaths); Belgium HR 1.84 (95% CI 1.29–2.63; P = 0.001; n = 245, 139 deaths) (Supplementary Table 4). All three cohort-specific hazard ratios were greater than one and statistically significant at the within-cohort level, consistent with a common positive association between %PV and mortality across centres. Notably, the largest cohort-specific effect was observed in Belgium, the cohort in which every patient received antifibrotic therapy. Because treatment status was invariant within this cohort, it cannot account for the association between %PV and mortality observed there.

This consistency was formally tested in the primary stratified Cox model fitted to the pooled patient-level data. A likelihood-ratio test comparing the primary model with an extended model that additionally included PV × cohort interaction terms yielded χ² = 3.24 on 2 degrees of freedom (P = 0.20), providing no evidence against the assumption of a common %PV effect across cohorts.

The robustness of the principal result to cross-cohort variation was further assessed by patient-level leave-one-out cross-validation. The multivariable stratified Cox model was re-fitted after sequentially excluding each cohort. When the Belgian cohort was excluded, the %PV hazard ratio was 1.37 (95% CI 1.15–1.63; P < 0.001; n = 537, 392 deaths); when the Turkish cohort was excluded, it was 1.43 (95% CI 1.17–1.75; P < 0.001; n = 484, 332 deaths); and when the Netherlands cohort was excluded, it was 1.54 (95% CI 1.25–1.89; P < 0.001; n = 543, 338 deaths). The pooled hazard ratio remained statistically significant in every iteration, and the relative ordering — smallest effect when Belgium was excluded, largest when the Netherlands was excluded — is consistent with the cohort-specific point estimates. Together, these analyses indicate that the prognostic association between higher %PV and increased all-cause mortality is robust to between-cohort variation and is not driven by any single cohort.

**Supplementary Note 3. Proportional hazards assumption and time-varying remediation**

The proportional hazards (PH) assumption of the primary multivariable stratified Cox model was assessed using scaled Schoenfeld residuals (cox.zph in the survival R package). The assumption was satisfied for the principal biomarker %PV (χ² = 0.02, P = 0.88), as well as for age (P = 0.53), sex (P = 0.59), smoking history (P = 0.63), and DLCO (P = 0.11). Violations were detected for antifibrotic therapy exposure (χ² = 9.38, P = 0.002) and FVC (χ² = 6.47, P = 0.011), with the global Schoenfeld test also indicating departure from proportionality across the full covariate set (χ² = 18.9, df = 7, P = 0.008).

In line with the pre-specified remediation strategy, a sensitivity analysis was performed in which time-varying coefficients were added for the two covariates that violated the assumption. Specifically, the multivariable stratified Cox model was re-fitted with tt(antifibrotic) and tt(FVC) terms, using tt(x) = x × log(time + 1) as the time transformation (as implemented via the tt argument of coxph). In this remediation model, the time-varying components were themselves statistically significant — tt(antifibrotic) HR 1.90 (95% CI 1.27–2.84; P = 0.002) and tt(FVC) HR 1.011 (95% CI 1.001–1.021; P = 0.031) — confirming that the protective effects of both antifibrotic therapy and higher baseline FVC attenuate with longer follow-up, consistent with previously reported observations.

Critically, the hazard ratio for %PV in this remediation model was 1.44 (95% CI 1.23–1.69; P < 0.001) — essentially identical to the principal estimate of 1.44 (95% CI 1.23–1.68; Supplementary Table 3). The prognostic association between baseline %PV and all-cause mortality is therefore robust to the proportional-hazards violations in other model components and does not depend on the assumption that the effects of antifibrotic therapy or FVC are constant over follow-up.

**Supplementary Table 1:** Baseline clinical characteristics of IPF patients and controls used for HiP-CT and molecular OMICs profiling.

|  | HiP-CT<br>(n=7) |  | snRNA-seq<br>(n=13) |  | 4i/GeoMx<br>(n=12) |  |
| --- | --- | --- | --- | --- | --- | --- |
|  | Control<br>(n=3) | IPF<br>(n=4) | Control<br>(n=7) | IPF<br>(n=6) | Control<br>(n=4) | IPF<br>(n=8) |
| **Age, years**<br>(mean ± SD) | 57.7 ± 32.1 | 59.0 ± 10.6 | 63.7 ± 2.8 | 58.7 ± 4.1 | 65.8 ± 4.5 | 56.9 ± 7.1 |
| **Male sex, n (%)** | 2 (66.7) | 2 (50.0) | 4 (57.1) | 4 (66.7) | 3 (75.0) | 5 (62.5) |
| **Ever smoker, n (%)** | n/a | 1 (25.0) | 4 (57.1) | 4 (66.7) | 2 (50.0) | 5 (62.5) |
| **Antifibrotic therapy, n (%)** | n/a | 2 (50.0) | n/a | 5 (83.3) | n/a | 6 (75.0) |
| **BMI**<br>(mean ± SD) | n/a | 26.7 ± 4.4 | 25.3 ± 3.4 | 25.3 ± 2.2 | 29.0 ± 3.8 | 24.4 ± 2.7 |
| **FVC, % predicted**<br>(mean ± SD) | n/a | 45.8 ± 14.2 | 97.6 ± 16.3 | 41.5 ± 15.8 | 121.3 ± 15.5 | 39.5 ± 14.3 |
| **FEV1, % predicted**<br>(mean ± SD) | n/a | 48.5 ± 17.2 | 96.1 ± 16.3 | 47.7 ± 16.7 | 118.8 ± 18.4 | 45.5 ± 15.0 |
| **FEV1 / FVC, %**<br>(mean ± SD) | n/a | 83.8 ± 6.9 | 77.6 ± 5.2 | 92.7 ± 6.8 | 75.3 ± 2.4 | 93.0 ± 6.6 |
| **TLC, % predicted**<br>(mean ± SD) | n/a | 52.3 ± 8.6 | 99.9 ± 8.8 | 52.0 ± 8.4 | 108.0 ± 13.7 | 51.1 ± 7.4 |
| **RV, % predicted**<br>(mean ± SD) | n/a | 75.5 ± 17.5 | 106.3 ± 14.5 | 56.8 ± 11.6 | 96.0 ± 12.9 | 60.8 ± 17.8 |

Footnote: Continuous variables are presented as mean ± SD. FVC, forced vital capacity. FEV1, forced expiratory volume in 1 second. TLC, total lung capacity. RV, residual volume.

**Supplementary table 2.**
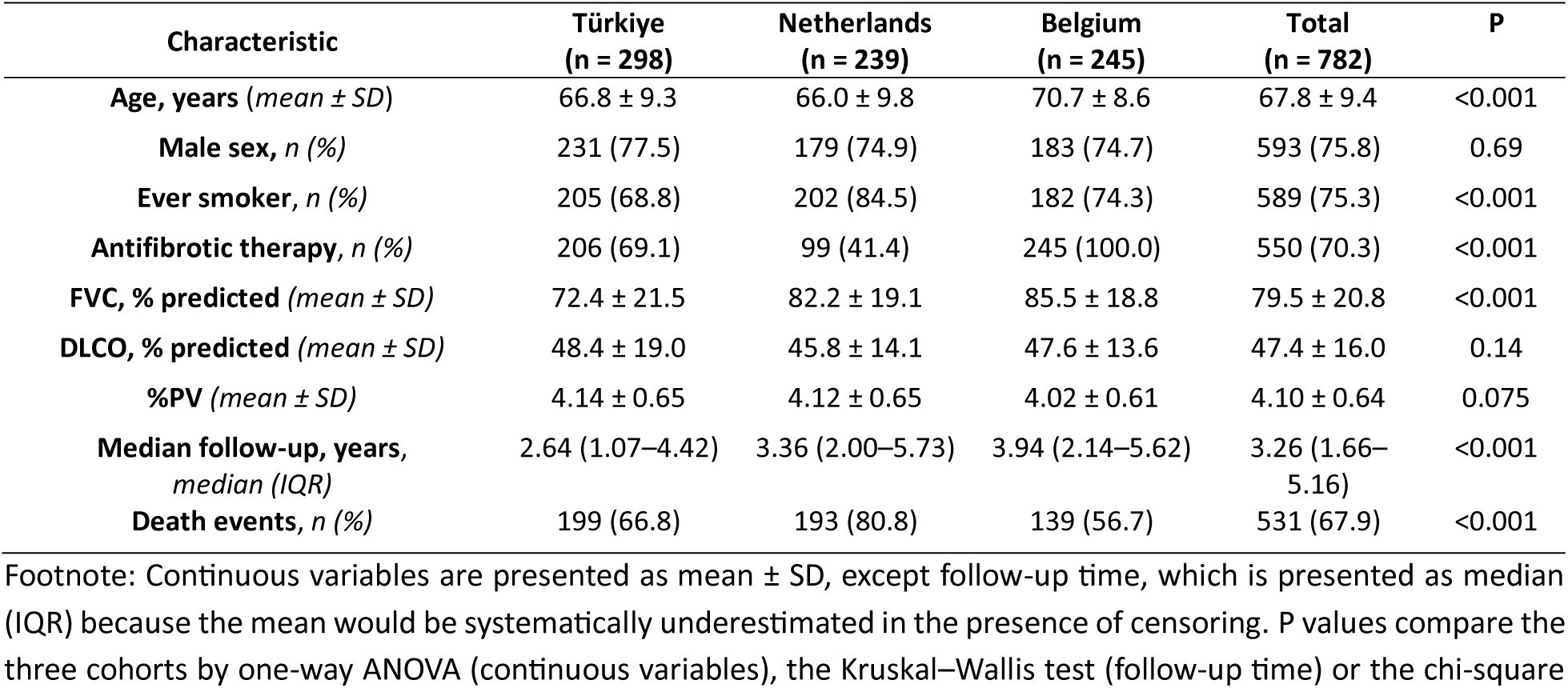
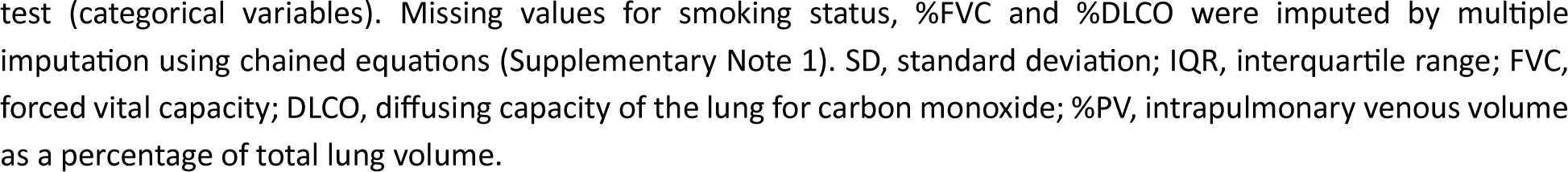
Baseline clinical characteristics of IPF patients across three clinical CT cohorts.

| Characteristic | Türkiye<br>(n = 298) | Netherlands<br>(n = 239) | Belgium<br>(n = 245) | Total<br>(n = 782) | P |
| --- | --- | --- | --- | --- | --- |
| **Age, years (mean ± SD)** | 66.8 ± 9.3 | 66.0 ± 9.8 | 70.7 ± 8.6 | 67.8 ± 9.4 | <0.001 |
| **Male sex, n (%)** | 231 (77.5) | 179 (74.9) | 183 (74.7) | 593 (75.8) | 0.69 |
| **Ever smoker, n (%)** | 205 (68.8) | 202 (84.5) | 182 (74.3) | 589 (75.3) | <0.001 |
| **Antifibrotic therapy, n (%)** | 206 (69.1) | 99 (41.4) | 245 (100.0) | 550 (70.3) | <0.001 |
| **FVC, % predicted (mean ± SD)** | 72.4 ± 21.5 | 82.2 ± 19.1 | 85.5 ± 18.8 | 79.5 ± 20.8 | <0.001 |
| **DLCO, % predicted (mean ± SD)** | 48.4 ± 19.0 | 45.8 ± 14.1 | 47.6 ± 13.6 | 47.4 ± 16.0 | 0.14 |
| **%PV (mean ± SD)** | 4.14 ± 0.65 | 4.12 ± 0.65 | 4.02 ± 0.61 | 4.10 ± 0.64 | 0.075 |
| **Median follow-up, years, median (IQR)** | 2.64 (1.07–4.42) | 3.36 (2.00–5.73) | 3.94 (2.14–5.62) | 3.26 (1.66–5.16) | <0.001 |
| **Death events, n (%)** | 199 (66.8) | 193 (80.8) | 139 (56.7) | 531 (67.9) | <0.001 |

Footnote: Continuous variables are presented as mean ± SD, except follow-up time, which is presented as median (IQR) because the mean would be systematically underestimated in the presence of censoring. P values compare the three cohorts by one-way ANOVA (continuous variables), the Kruskal–Wallis test (follow-up time) or the chi-square
test (categorical variables). Missing values for smoking status, %FVC and %DLCO were imputed by multiple imputation using chained equations (Supplementary Note 1). SD, standard deviation; IQR, interquartile range; FVC, forced vital capacity; DLCO, diffusing capacity of the lung for carbon monoxide; %PV, intrapulmonary venous volume as a percentage of total lung volume.

**Supplementary table 3.** Stratified Cox Regression Analysis for Pulmonary Vein Volume (%PV).

| Variable | Univariate HR (95% CI) | P | Multivariable HR (95% CI) | P |
| --- | --- | --- | --- | --- |
| **%PV** (per 1-unit) | 1.57 (1.38–1.79) | <0.001 | 1.44 (1.23–1.68) | <0.001 |
| **Age, years** (per year) | 1.02 (1.01–1.03) | <0.001 | 1.02 (1.01–1.04) | <0.001 |
| **Male sex** (vs female) | 1.51 (1.21–1.87) | <0.001 | 1.53 (1.18–1.97) | 0.001 |
| **Ever smoker** (vs never) | 1.31 (1.06–1.61) | 0.013 | 1.06 (0.83–1.36) | 0.635 |
| **Antifibrotic therapy** (vs none) | 0.63 (0.50–0.79) | <0.001 | 0.62 (0.49–0.79) | <0.001 |
| **FVC, % predicted** (per 1%) | 0.98 (0.97–0.98) | <0.001 | 0.99 (0.98–1.00) | <0.001 |
| **DLCO, % predicted** (per 1%) | 0.96 (0.95–0.97) | <0.001 | 0.96 (0.96–0.97) | <0.001 |

Hazard ratios from stratified Cox proportional-hazards regression with cohort as the stratification variable. Univariate column: each row from a separate model containing that variable and the stratification variable only. Multivariable column: all seven variables fitted simultaneously. Lung-transplant recipients were censored at the date of transplantation. Missing values for smoking status, %FVC and %DLCO were imputed by multiple imputation using chained equations (Supplementary Note 1). CI, confidence interval; HR, hazard ratio; FVC, forced vital capacity; DLCO, diffusing capacity for carbon monoxide.

**Supplementary table 4.** Per-cohort multivariable Cox proportional-hazards regression for all-cause mortality.

| Variable | Türkiye (n=298) |  | Netherlands (n=239) |  | Belgium (n=245) |  |
| --- | --- | --- | --- | --- | --- | --- |
|  | HR (95% CI) | P | HR (95% CI) | P | HR (95% CI) | P |
| **%PV** (per 1-unit) | 1.40 (1.09–1.79) | 0.009 | 1.32 (1.02–1.72) | 0.037 | 1.84 (1.29–2.63) | 0.001 |
| **Age** (per year) | 1.02 (1.00–1.03) | 0.071 | 1.02 (1.00–1.04) | 0.119 | 1.04 (1.02–1.07) | <0.001 |
| **Male sex** (vs female) | 1.08 (0.64–1.83) | 0.765 | 1.55 (1.02–2.35) | 0.038 | 1.79 (1.09–2.96) | 0.023 |
| **Ever smoker** (vs never) | 1.39 (0.87–2.23) | 0.166 | 1.01 (0.64–1.58) | 0.980 | 1.10 (0.71–1.71) | 0.674 |
| **Antifibrotic therapy** (vs none) | 0.38 (0.27–0.52) | <0.001 | 0.92 (0.68–1.26) | 0.606 | NE | NE |
| **FVC, % predicted** (per 1%) | 0.98 (0.97–0.99) | 0.001 | 0.99 (0.98–1.01) | 0.364 | 0.99 (0.98–1.00) | 0.079 |
| **DLCO, % predicted** (per 1%) | 0.97 (0.96–0.98) | <0.001 | 0.96 (0.95–0.97) | <0.001 | 0.95 (0.93–0.97) | <0.001 |

Multivariable Cox proportional-hazards models were fitted separately within each cohort. HR, hazard ratio; CI, confidence interval; FVC, forced vital capacity; DLCO, diffusing capacity of the lung for carbon monoxide; NE, not estimable. Antifibrotic therapy could not be estimated in the Belgian cohort because all patients had received antifibrotic therapy. Missing values for smoking status, %FVC and %DLCO were imputed by multiple imputation using chained equations (Supplementary Note 1). %PV is expressed per one-percentage-point increase.

**Supplementary table 5.** Primary and secondary antibodies for 4i staining.

| Primary antibody | Manufacturer | Lot number | Primary antibody dilution | Secondary antibody | Lot number | Secondary antibody dilution | Biological process |
| --- | --- | --- | --- | --- | --- | --- | --- |
| CLDN5 | Santa Cruz Biotechnology | sc-374221 | 1:100 | Donkey anti-mouse Alexa Fluor 488 | A21202 | 1:250 | Pan-Endothelial cells |
| TNC | Abcam | AB108930 | 1:100 | Donkey anti-rabbit Alexa Fluor 568 | A10042 | 1:250 | ECM, Myofibs |
| PDPN | R&D Systems | AF3670 | 1:100 | Donkey anti-sheep Alexa Fluor 647 | A21448 | 1:250 | AT1, LEC |
| E-cadherin | BD Biosciences | 610182 | 1:100 | Donkey anti-mouse Alexa Fluor 488 | A21202 | 1:250 | Pan epithelial cell marker |
| PLVAP | Novus Biologicals | NBP1-83911 | 1:300 | Donkey anti-rabbit Alexa Fluor 568 | A10042 | 1:250 | Peribronchiolar ECs |
| PTPRC | LSBio | LS-B14248 | 1:300 | Donkey anti-goat Alexa Fluor 647 | A21447 | 1:250 | Pan-leukocyte |
| aSMA | Sigma-Aldrich | A5228 | 1:1500 | Donkey anti-mouse Alexa Fluor 488 | A21202 | 1:250 | SMC and myofibroblasts |
| VWA1 | proteintech | 14322-1-AP | 1:300 | Donkey anti-rabbit Alexa Fluor 568 | A10042 | 1:250 | ectopic EC |
| SPP1 | R&D Systems | AF1433 | 1:200 | Donkey anti-goat Alexa Fluor 647 | A21447 | 1:250 | Macrophages/ ECM |
| CD20 | Abcam | AB9475 | 1:200 | Donkey anti-mouse Alexa Fluor 488 | A21202 | 1:250 | B cells |
| PRX | Sigma-Aldrich | HPA001868 | 1:500 | Donkey anti-rabbit Alexa Fluor 568 | A10042 | 1:250 | Capillaries |
| RAGE | R&D Systems | AF1145 | 1:100 | Donkey anti-goat Alexa Fluor 647 | A21447 | 1:250 | AT1, LEC |
| CD68 | Thermo Fisher Scientific | 14-0688-82 | 1:400 | Donkey anti-mouse Alexa Fluor 488 | A21202 | 1:250 | Pan-Macrophage |
| CD31 | abcam | AB28364 | 1:50 | Donkey anti-rabbit Alexa Fluor 568 | A10042 | 1:250 | pan EC |
| CD206 | R&D Systems | AF2534 | 1:50 | Donkey anti-goat Alexa Fluor 647 | A21447 | 1:250 | M2 macrophages (anti-inflammatory, pro-fibrosis) |
| KRT8 | DSHB | AB_531826 | 1:200 | Donkey anti-rat Alexa Fluor 488 | A21208 | 1:250 | Krt8-ADI, Bronchial epithelial cells |
| KRT17 | Sigma-Aldrich | HPA000453 | 1:150 | Donkey anti-rabbit Alexa Fluor 568 | A10042 | 1:250 | Basal cells |
| KRT5 | Biolegend | 905903 | 1:800 | Donkey anti-chicken Alexa Fluor 647 | 703-605-155 | 1:250 | Basal cells |
| CD3 | Abcam | AB11089 | 1:100 | Donkey anti-rat Alexa Fluor 488 | A21208 | 1:250 | T cells |
| SFTPC | Sigma-Aldrich | HPA010928 | 1:200 | Donkey anti-rabbit Alexa Fluor 568 | A10042 | 1:250 | AT2 |
| CD90 | R&D Systems | AF2067 | 1:50 | Donkey anti-sheep Alexa Fluor 647 | A21448 | 1:250 | Fibroblasts |
| SCGB1A1 | R&D Systems | MAB4218 | 1:100 | Donkey anti-rat Alexa Fluor 488 | A21208 | 1:250 | Goblet /Club cells |
| MZB1 | Sigma-Aldrich | HPA043745 | 1:2000 | Donkey anti-rabbit Alexa Fluor 568 | A10042 | 1:250 | Plasma cells (CD45- or low) |
| POSTN | Cell Signaling Technology | 20302 | 1:200 | Donkey anti-rabbit Alexa Fluor 568 | A10042 | 1:250 | ECM/Myofibs |

**Supplementary Figure 1.**
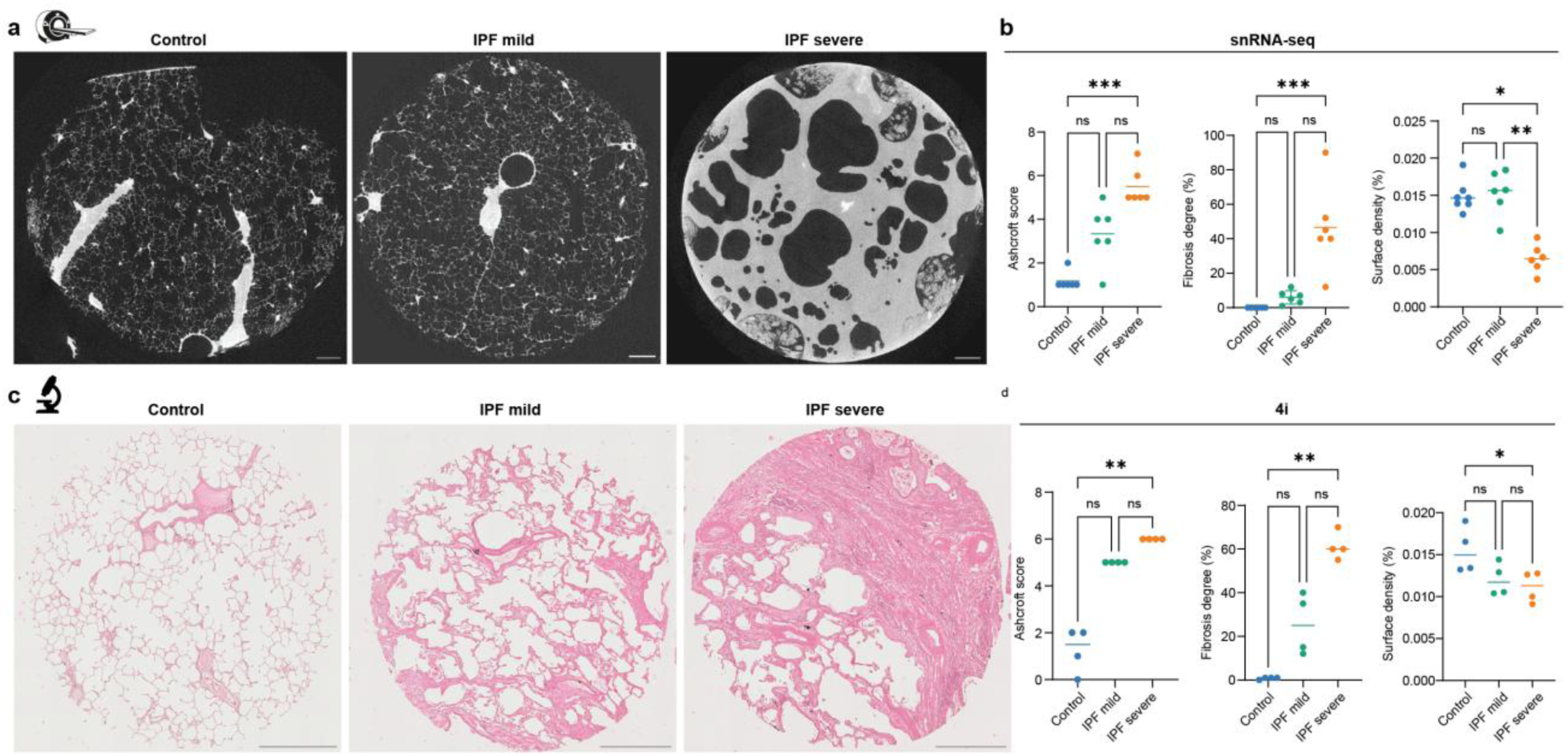
Micro-CT and histopathology-based staging of IPF severity. (a) Representative micro-computed tomography (micro-CT) images from control, mild IPF, and severe IPF samples. Scale bar, 1 mm. (b) Dot plots showing patient-level measurements of Ashcroft score (left), fibrosis degree (middle), and surface density (right) across IPF severity in the snRNA-seq dataset. (c) Representative hematoxylin and eosin (H&E) images from control, mild IPF, and severe IPF samples. Scale bar, 1 mm. (d) Dot plots showing patient-level measurements of Ashcroft score (left), fibrosis degree (middle), and surface density (right) across IPF severity. A comparison among the three groups was performed using the Kruskal–Wallis test, followed by Dunn’s post hoc test for pairwise comparisons.

**Supplementary Figure 2.**
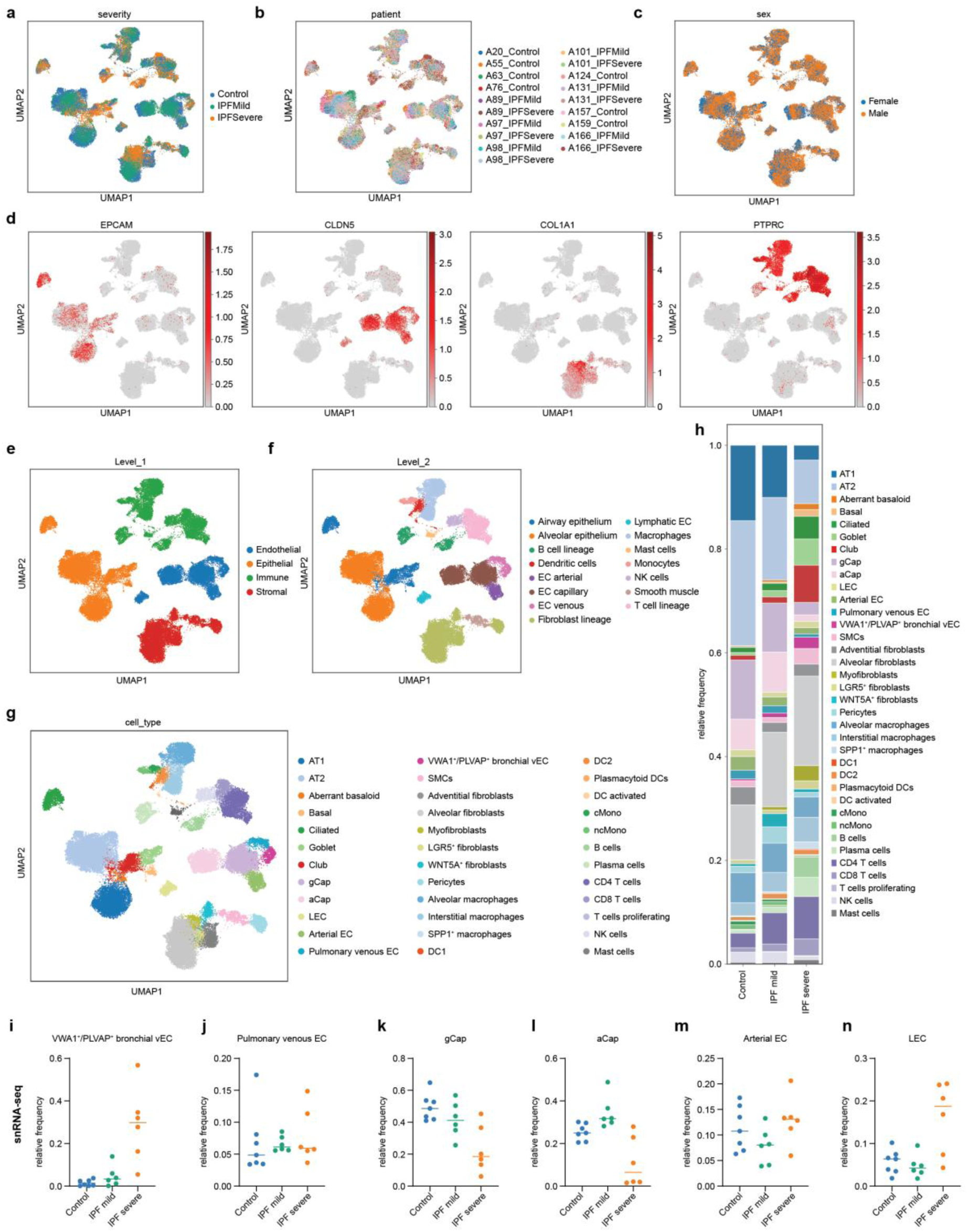
Overview of the snRNA-seq dataset (63,072 cells). (a–c) Uniform Manifold Approximation and Projection (UMAP) of the entire dataset coloured by fibrosis severity (a), individual donor (b) and sex (c). (d) UMAP showing the expression of canonical lineage markers used for major compartment annotation (EPCAM, epithelial cells; CLDN5, endothelial cells; COL1A1, mesenchymal cells; and PTPRC, immune cells). (e–g) UMAPs showing the major cellular compartment annotation (e), broad cell type annotation based on the Human Lung Cell Atlas (HLCA) hierarchy (Level 2; 15 cell types; f), and fine-grained cell type/state annotation (36 cell types/states; g). (h) Stacked bar plots showing the relative cellular composition of the entire dataset across control, mild IPF and severe IPF samples. (i–n) Dot plots showing changes in the relative frequencies of the different endothelial cell populations across control, mild IPF and severe IPF. The line indicates the mean.

**Supplementary Figure 3.**
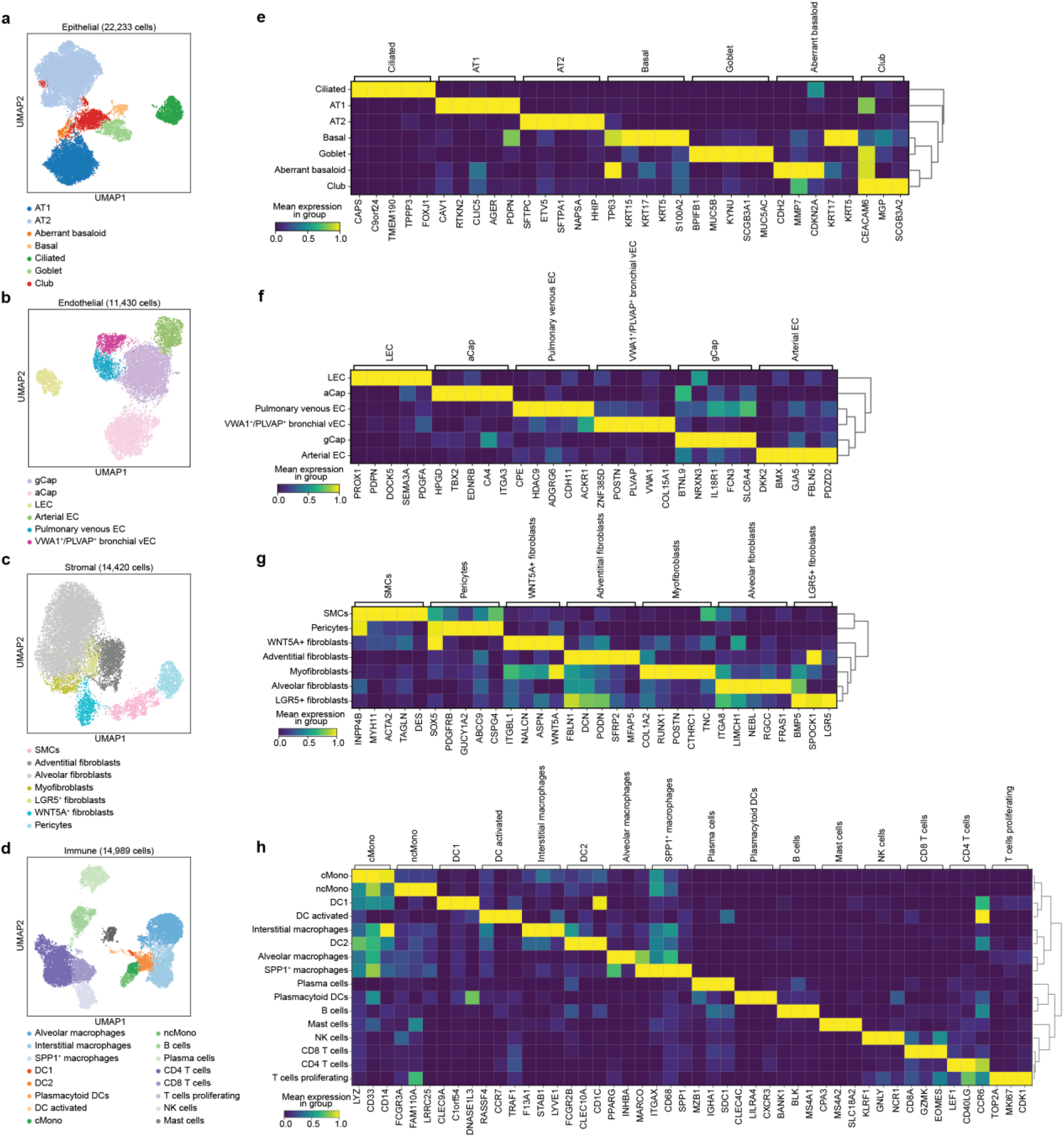
Overview of the major cellular compartments identified by snRNA-seq and their representative marker genes. (a–d) Uniform Manifold Approximation and Projection (UMAP) of the epithelial (a, 22,233 cells), endothelial (b, 11,430 cells), stromal (c, 14,420 cells) and immune (d, 14,989 cells) cell compartments. (e–h) Matrix plots showing representative marker genes for each major cellular compartment: epithelial (e), endothelial (f), mesenchymal (g) and immune (h) cells. Colour intensity indicates the scaled mean gene expression.

**Supplementary Figure 4.**
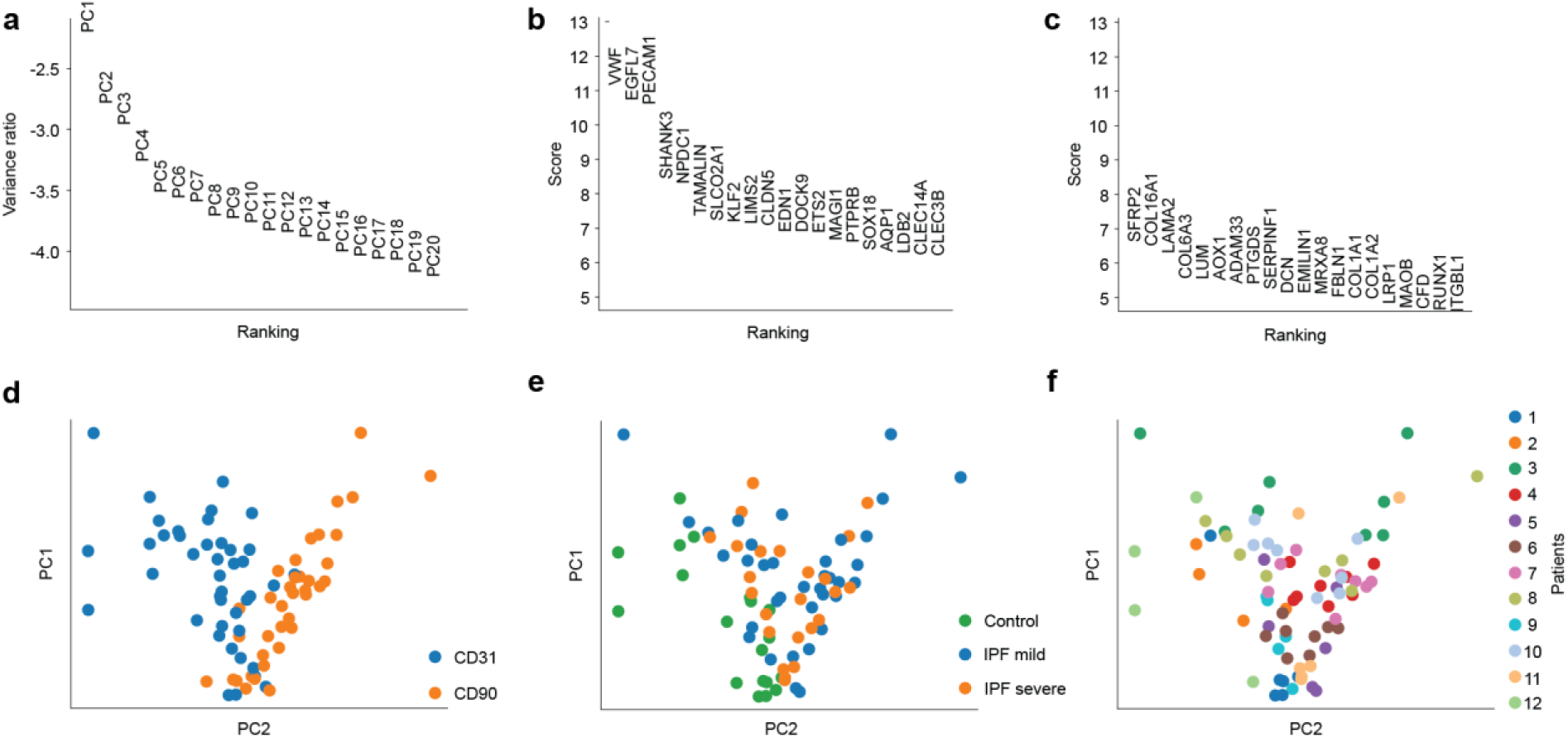
Principal component analysis of GeoMx CD31⁺ and CD90⁺ regions of interest. (a) Explained-variance ratios for the first 20 principal components, showing the proportion of transcriptional variation captured by each component. (b–c) Ranked compartment-associated genes in the CD31⁺ (b) and CD90⁺ (c) segments. Enrichment of canonical vascular/endothelial transcripts in CD31⁺ regions and stromal/mesenchymal transcripts in CD90⁺ regions support the specificity of the segmentation strategy. (d–f) PCA projections colored by segmentation compartment (d), disease severity (e), and patient identity (f), illustrating the contributions of tissue compartment, disease state and interpatient variability to global transcriptional differences.

**Supplementary Figure 5.**
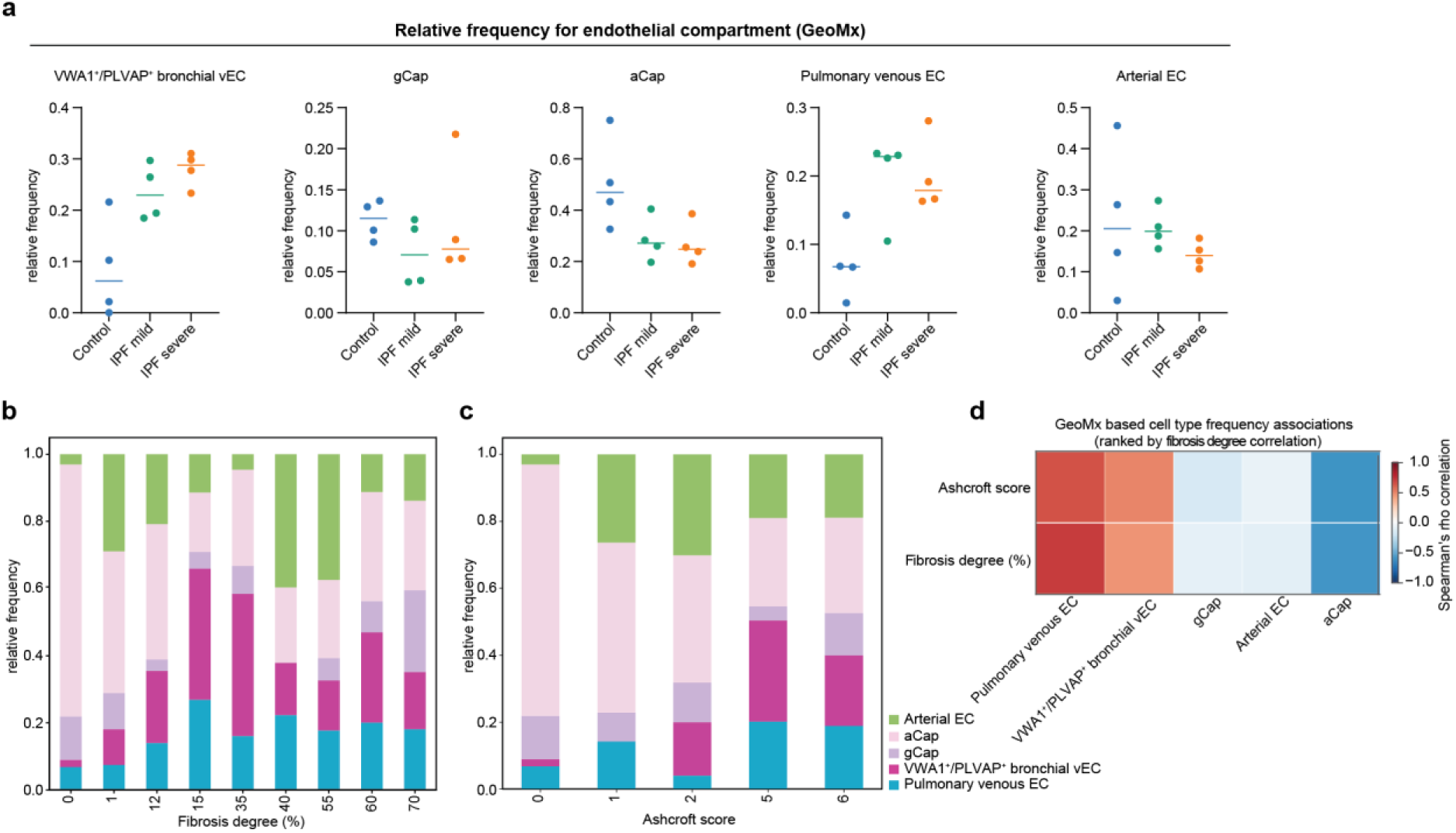
Endothelial cell abundance across IPF severity in the GeoMx dataset. (a) Dot plot showing the relative frequency of deconvoluted endothelial cell (EC) subtypes across different stages of IPF severity. The line indicates the mean. (b–c) Stacked bar plots showing the relative frequency of deconvoluted EC subtypes stratified by fibrosis degree (b) and Ashcroft score (c). (d) Heatmap showing Spearman correlation coefficients between endothelial cell population frequencies and fibrosis degree or Ashcroft score. Endothelial cell populations are ranked according to their correlation with fibrosis degree.

**Supplementary Figure 6.**
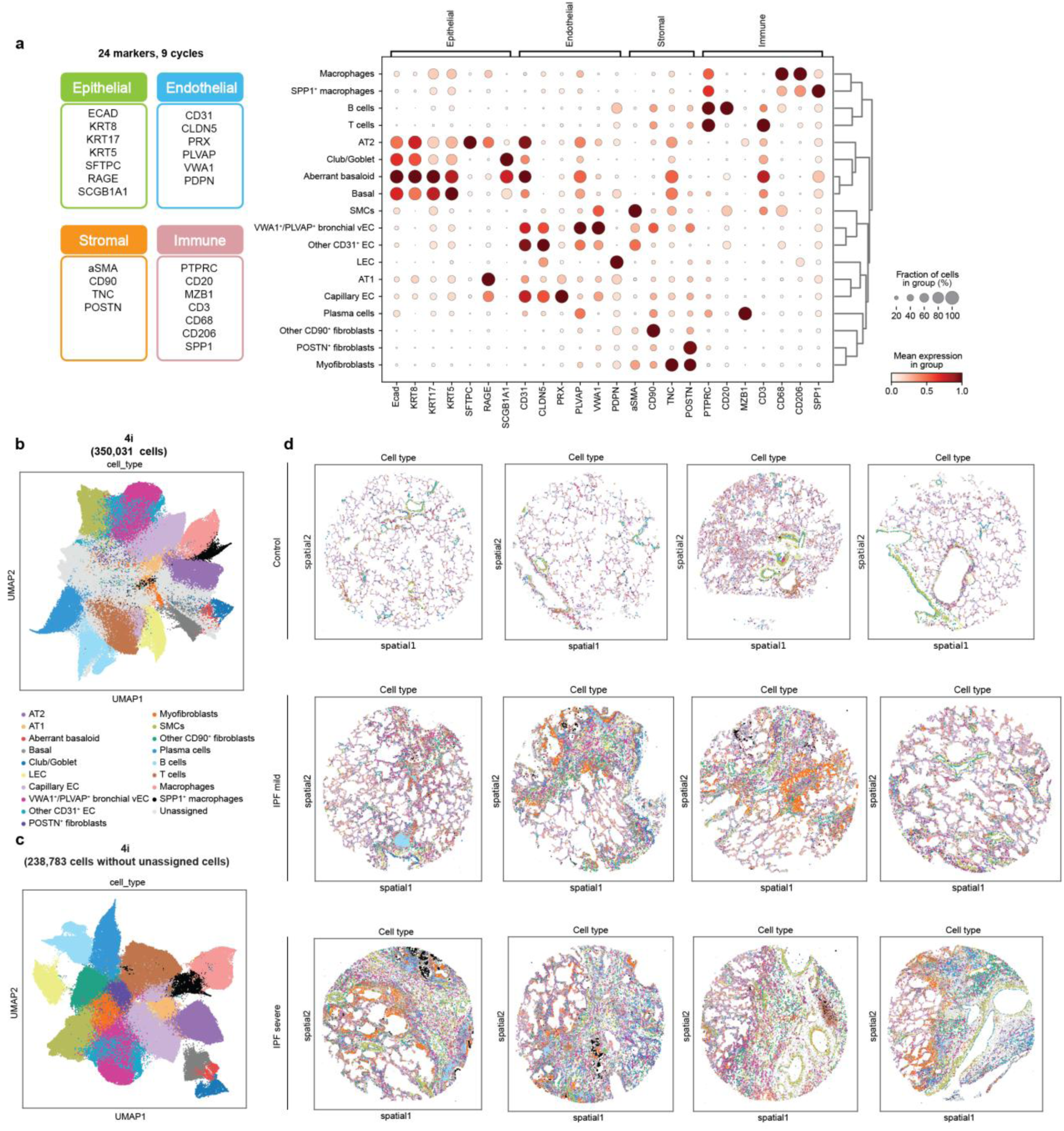
Spatial profiling of the tissue microarray (TMA). (a) Overview of the 4i experimental design comprising 24 protein markers measured across 9 imaging cycles. Dot plots summarize marker protein expression across all annotated cell populations within the epithelial, endothelial, stromal, and immune compartments. Colour intensity indicates the scaled mean gene expression. (b–c) UMAP visualization of all cells (n = 350,031 cells; b) and of annotated cell populations after excluding unassigned cells (n = 238,783 cells; c) in the 4i dataset. (d) Spatial UMAP representation of cell populations for each individual lung TMA sample.

**Supplementary Figure 7.**
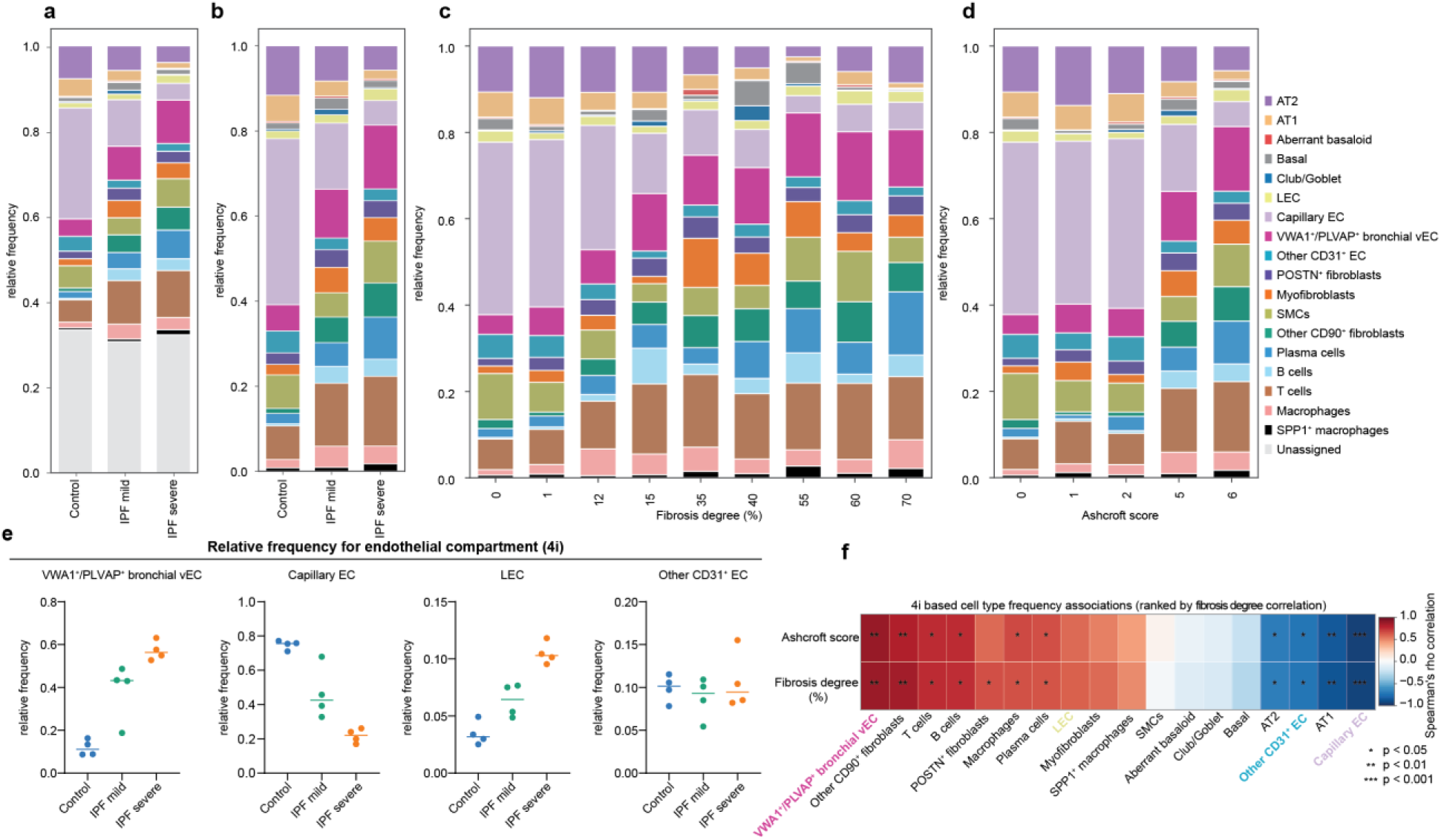
Cellular composition across IPF severity in the 4i dataset. (a-b) Relative frequency of all cell populations (a) and annotated cell populations after excluding unassigned cells (b) across different stages of IPF severity. (c–d) Relative frequencies of cell populations stratified by fibrosis degree (c) and Ashcroft score (d). (e) Dot plot showing the relative frequency of endothelial cells across IPF severity. The line indicates the mean. (f) Heatmap showing FDR-corrected Spearman correlation coefficients between cell population frequencies and fibrosis degree or Ashcroft score. Cell populations are ranked according to their correlation with fibrosis degree.

**Supplementary Figure 8.**
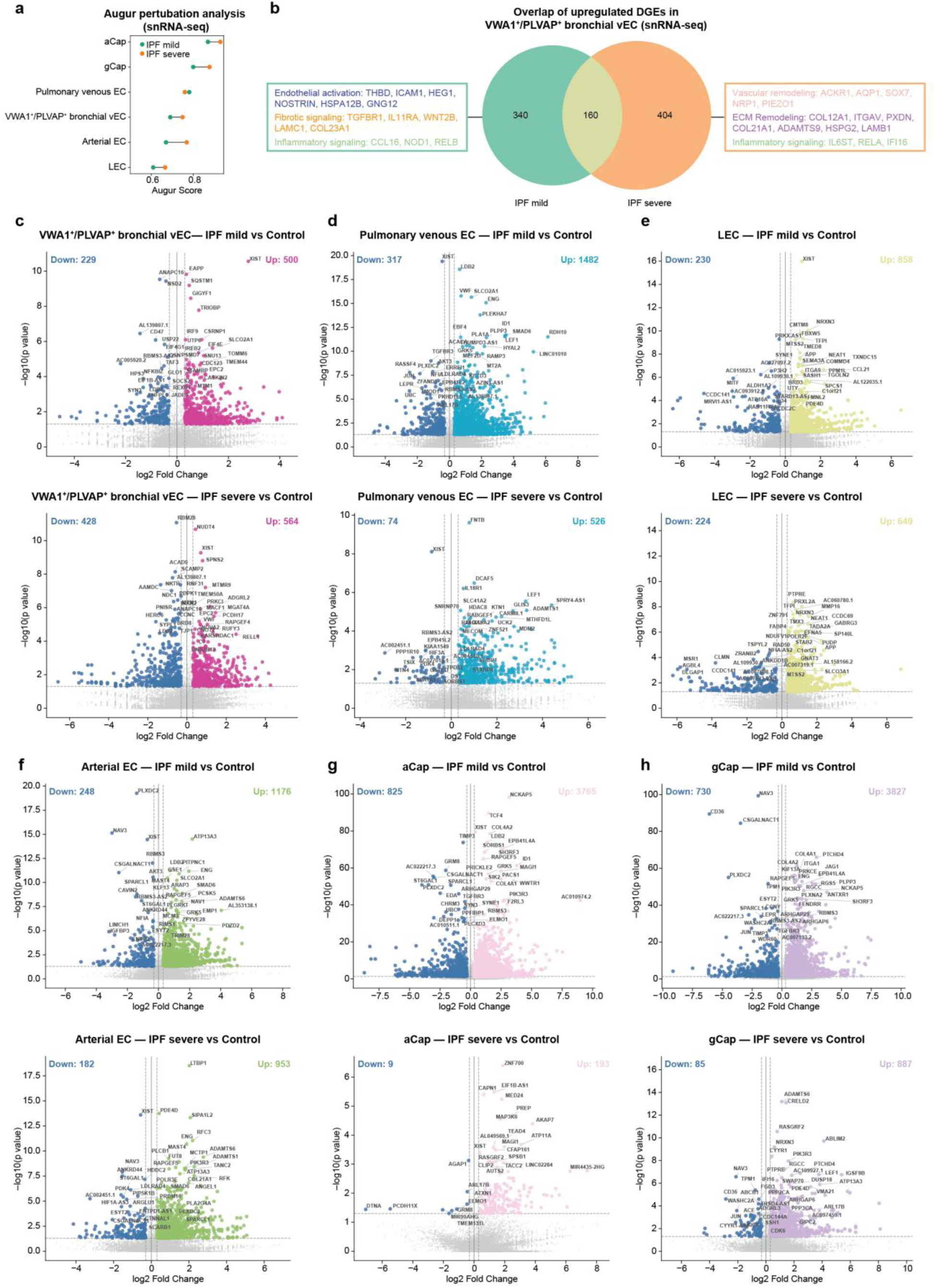
Endothelial cell type-specific transcriptional changes across IPF severity. (a) Endothelial cell type-resolved Augur perturbation analysis comparing mild IPF with control and severe IPF with control. (b) Venn diagram showing the gene pathways uniquely identified in IPF mild and IPF severe. (c–h) Volcano plots showing differentially expressed genes in VWA1⁺/PLVAP⁺ bronchial venous endothelial cells (ECs; c), pulmonary venous ECs (d), lymphatic ECs (LECs; e), arterial ECs (f), aerocytes (aCap; g) and general capillary ECs (gCap; h). For each endothelial cell type, the upper plot shows mild IPF versus control and the lower plot shows severe IPF versus control. Genes meeting the significance thresholds of an absolute log₂ fold change > 0.3 and raw *P* < 0.05 are highlighted, with the top 20 most significant genes labeled.

**Supplementary Figure 9.**
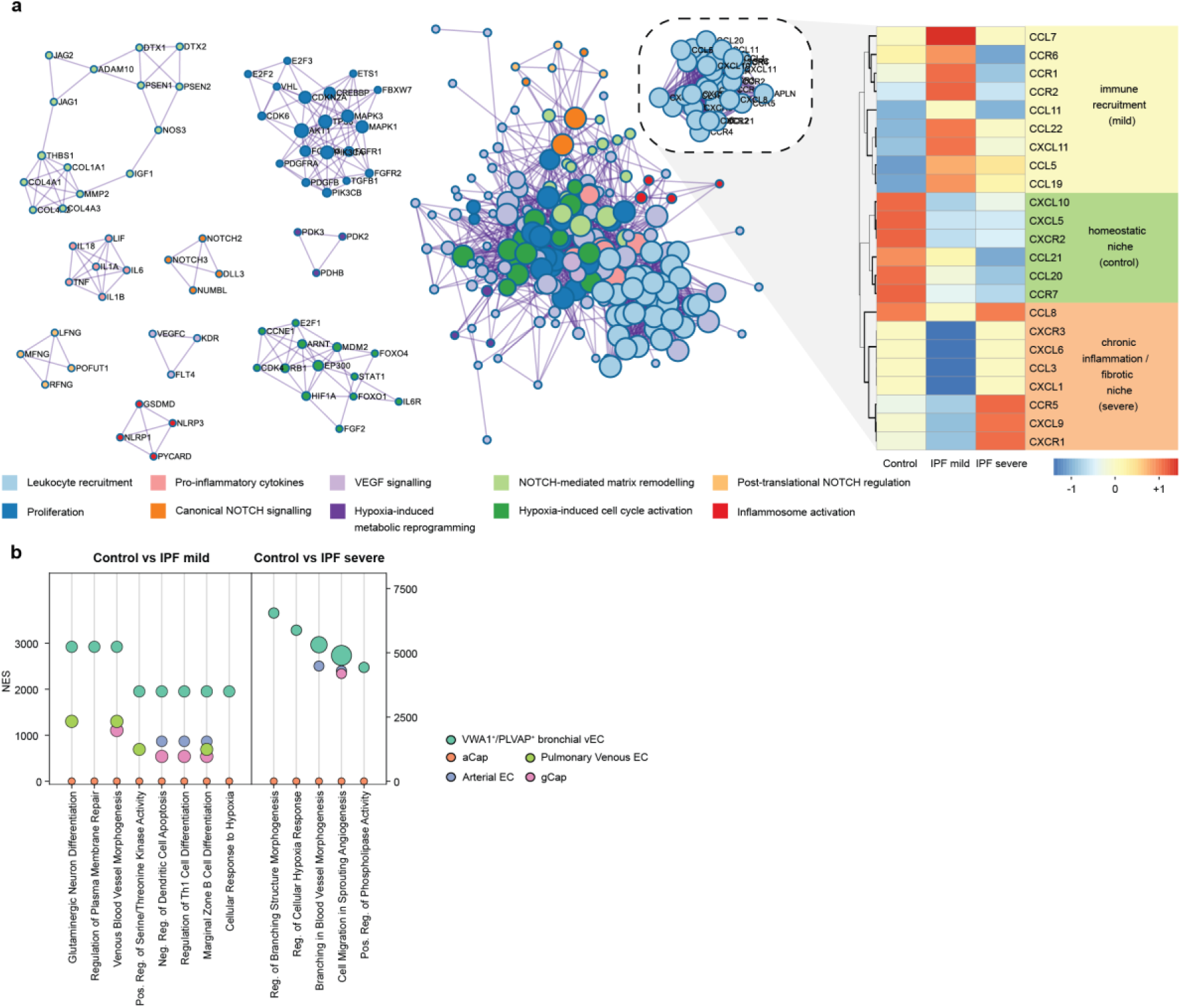
Transcriptional programs associated with disease severity in VWA1⁺/PLVAP⁺ bronchial venous endothelial cells in the GeoMx dataset. (a) Metascape network visualization of genes upregulated in VWA1⁺/PLVAP⁺ bronchial venous endothelial cells (ECs) from both mild and severe IPF regions relative to control lung regions. Genes encoding chemokines implicated in immune-cell recruitment are highlighted. The adjacent heatmap shows the relative expression of these chemokines and their corresponding receptors across control, mild-IPF, and severe-IPF regions. Expression values are standardized per gene and presented as z-scores. (b) Dot plot showing the top enriched pathways in VWA1⁺/PLVAP⁺ bronchial venous ECs for mild IPF versus control and severe IPF versus control, based on the Gene Ontology Biological Process 2025 gene-set library. Normalized enrichment scores (NES) indicate the direction and magnitude of pathway enrichment and dot size represents statistical significance as −log₁₀(FDR-adjusted P value).

**Supplementary Figure 10.**
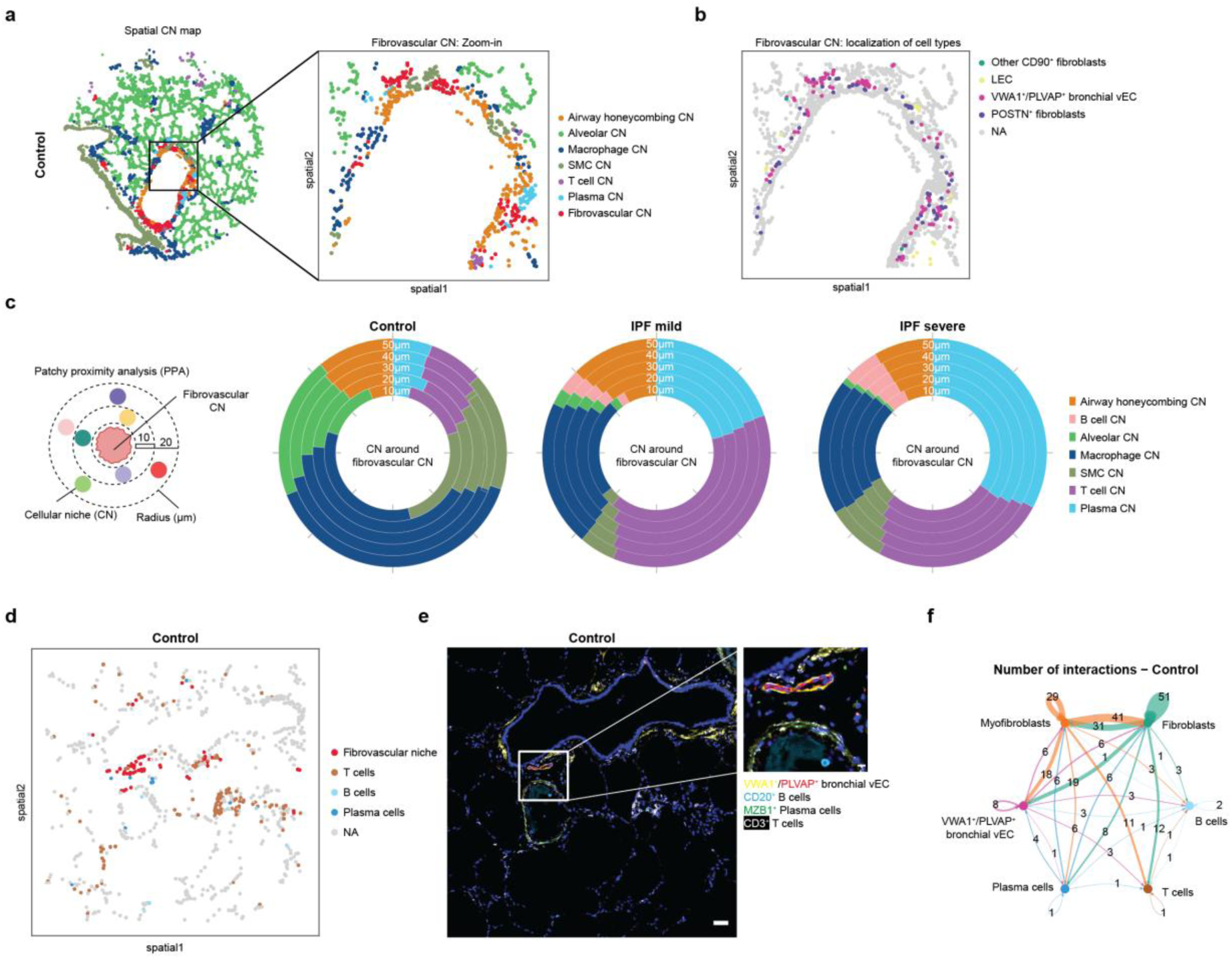
Changes in cellular niche (CN) composition surrounding the fibrovascular niche in controls and different IPF severities in the 4i dataset. (a) Representative spatial CN map of a control lung tissue sample. (b) Spatial localization of cell populations within the fibrovascular niche in a representative control sample. (c) Proximity profile analysis (PPA) showing changes in the cellular niche (CN) composition surrounding the fibrovascular niche across different stages of IPF severity. (d) Spatial UMAP showing the distribution of cell populations surrounding the fibrovascular niche in a representative control sample. (e) Corresponding immunofluorescence (IF) images highlighting cell populations surrounding the fibrovascular niche in a representative control sample. Scale bar, 50 µm (overview) and 10 µm (magnified views). (f) Circle plot illustrating the number of cell–cell interactions among cell populations around the fibrovascular niche in the control samples.

**Supplementary Figure 11.**
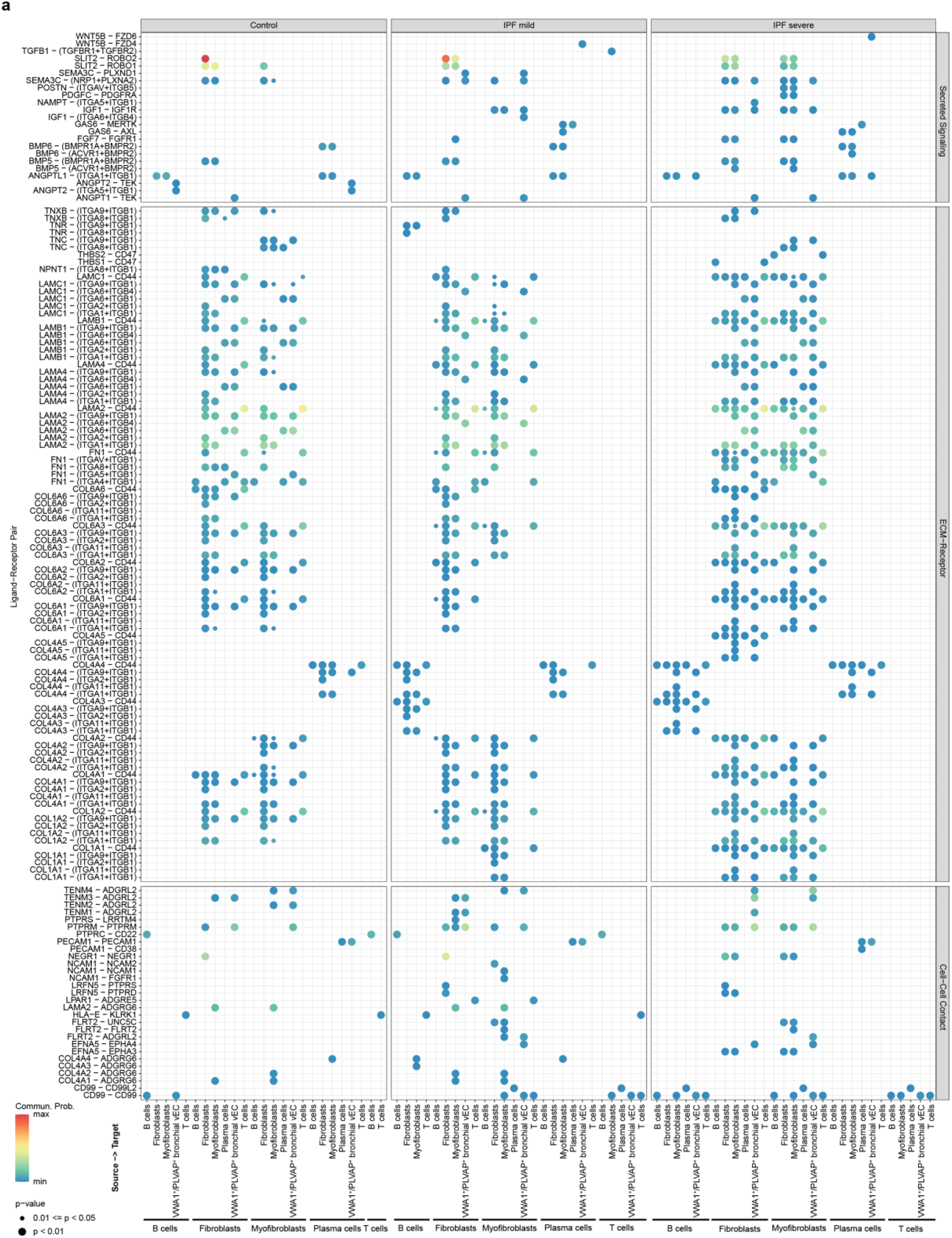
Cell-cell communication among cell populations surrounding fibrovascular niche in the snRNA-seq dataset. (a) Cell–cell communication network among cell populations surrounding the fibrovascular niche, inferred from the snRNA-seq dataset. Interactions with P > 0.05 were excluded. Dot size represents the level of statistical significance, with larger dots indicating lower P values. Dot color represents estimated communication probability, ranging from lower (blue) to higher (red) probability.

**Supplementary Figure 12.**
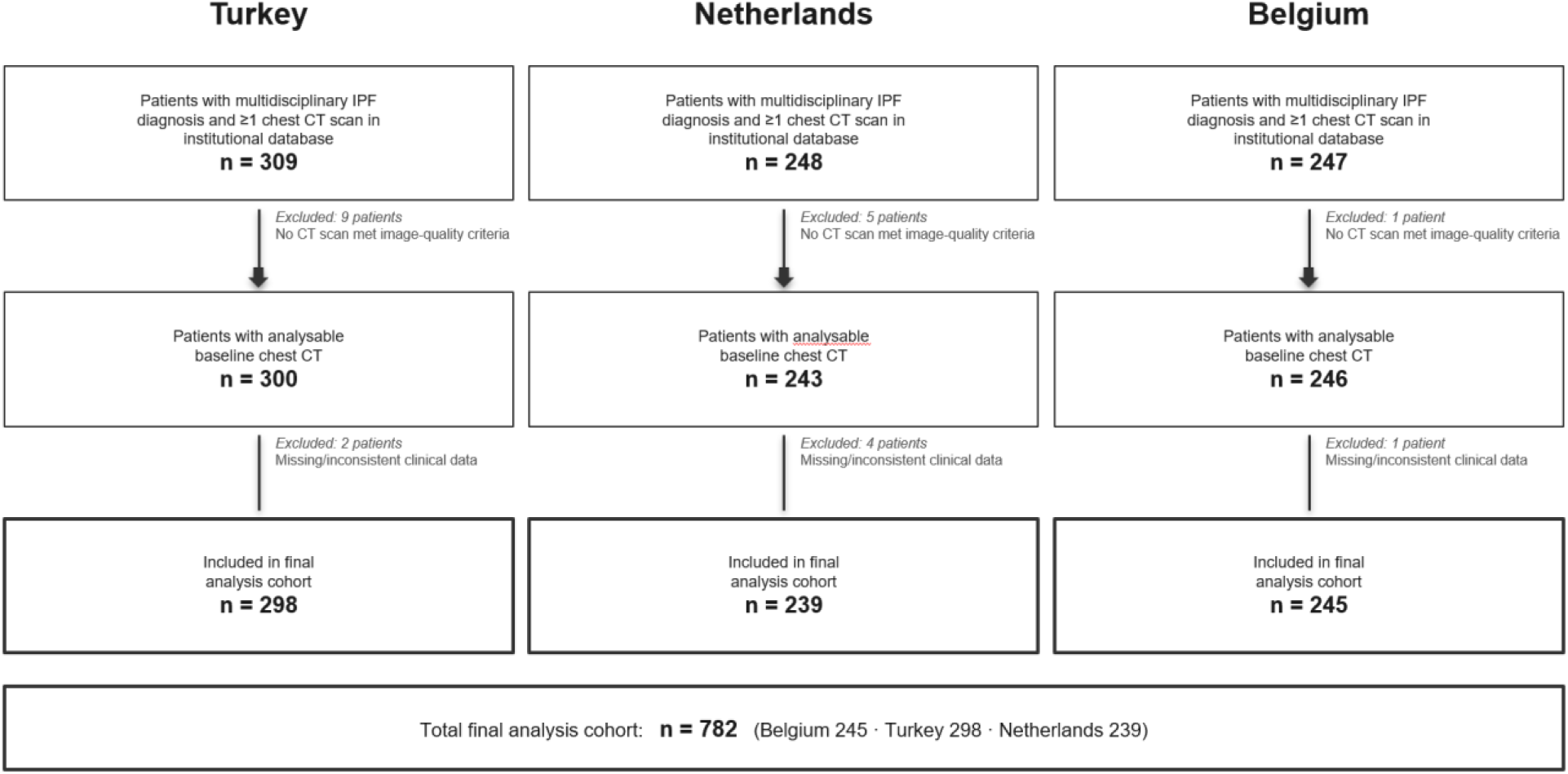
Patient selection for clinical CT analysis. Flow diagram of patient inclusion and exclusion across the three independent IPF cohorts contributing to the clinical CT survival analysis. The starting population at each participating centre comprised patients with a multidisciplinary diagnosis of IPF and at least one chest CT scan available in the institutional imaging database (Türkiye, n = 309; the Netherlands, n = 248; Belgium, n = 247). All available chest CT scans for each patient were quality screened and were rejected if they exhibited low z-axis resolution or excessive slice thickness, motion artefact, incomplete lung coverage, or coexisting major thoracic pathology preventing reliable vascular segmentation (e.g., pleural effusion, pneumothorax, prior lobectomy or lobar resection). For patients with multiple available scans, the earliest scan meeting image-quality criteria was retained as the baseline scan. Patients in whom none of the available CT scans met these criteria were excluded (Belgium, n = 1; Türkiye, n = 9; the Netherlands, n = 5). Patients with missing or internally inconsistent baseline clinical data, defined as missing or inconsistent records of age, sex, antifibrotic therapy exposure, or follow-up information,were additionally excluded (Belgium, n = 1; Türkiye, n = 2; the Netherlands, n = 4). The final analysis cohort comprised 782 patients ( Türkiye, n = 298; the Netherlands, n = 239; Belgium, n = 245).

**Supplementary Figure 13.**
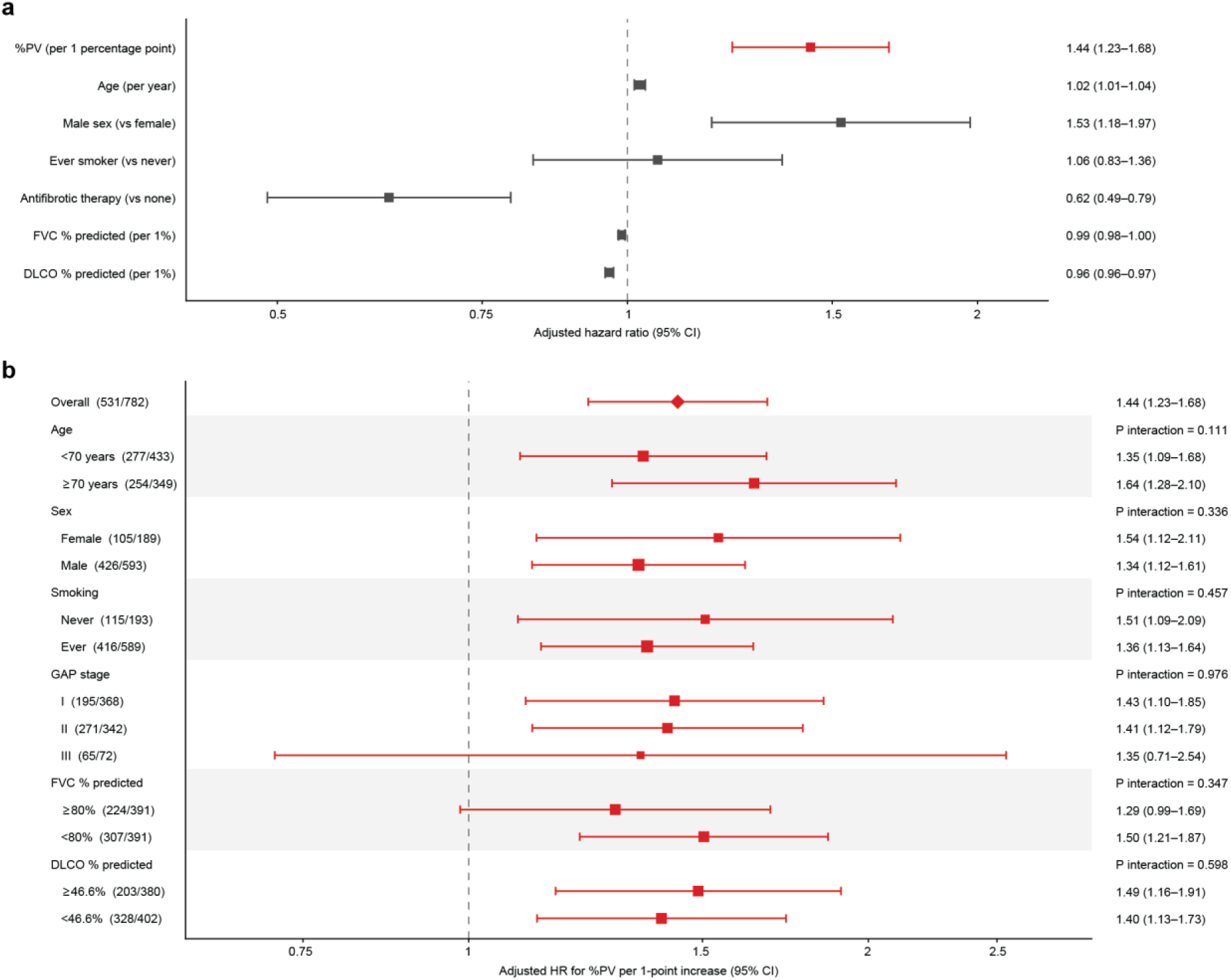
Multivariable and subgroup analyses of the association between %PV and all-cause mortality in the pooled clinical CT cohorts. (a) Forest plot of the primary multivariable Cox proportional-hazards model stratified by cohort (n = 782, 531 deaths), showing adjusted hazard ratios for all covariates. Continuous variables are expressed per one-unit increase (%PV per one percentage point, age per year, and %FVC and %DLCO per 1% predicted). (b) Adjusted hazard ratios for death per one-percentage-point increase in %PV within subgroups defined by age (<70 vs. ≥70 years), sex, smoking status, GAP stage (I, 0–3 points; II, 4–5 points; III, 6–8 points), %FVC (≥80% vs. <80% predicted) and %DLCO (split at the pooled median, 46.6% predicted). Within each subgroup, models were adjusted for age, sex, smoking status, antifibrotic therapy, %FVC and %DLCO, omitting the defining variable for sex and smoking status, and stratified by cohort. Numbers in brackets indicate deaths/total patients; for subgroups defined by imputed variables (GAP stage, %FVC and %DLCO), numbers are averaged across the 20 imputed datasets. P values for interaction were obtained by adding a %PV × subgroup interaction term to the model fitted to the full cohort; for GAP stage, the two interaction terms were tested jointly. The diamond indicates the overall estimate, error bars indicate 95% confidence intervals, and symbol size reflects subgroup size. Missing %FVC and %DLCO values were handled by multiple imputation (m = 20), and estimates were pooled using Rubin’s rules.

**Supplementary file 1. Differential expressed genes for endothelial cells.**

Mast DGE table includes all genes expressed in endothelial cells including aCap, gCap, arterial EC, LEC, VWA1+/PLVAP+ bronchial venous EC, pulmonary venous EC comparing IPF mild vs control, IPF severe vs control.

**Supplementary file 2. Upregulated GO pathways for endothelial cells.**

IPF_GO_pathway table contains upregulated pathways for all endothelial cells, including VWA1+/PLVAP+ bronchial venous EC, pulmonary venous EC, gCap, aCap, arterial EC and LEC in IPF mild and IPF severe. log2FC>0.3, adjusted P value<0.2.

## Notes

https://doi.org/10.5281/zenodo.22685651

https://github.com/gote-schniering-lab/2026_Yamada_Wang_et_al_Multiscale_mapping_of_venous_remodelling_in_IPF

### Summary of Updates

The clinical CT section has been extended with cohort-specific and pooled adjusted estimates, a dose-response analysis and subgroup analyses (revised Figure 7 and new Supplementary Figure 13), with corresponding updates to the Results, Methods and figure legends. The GEO accession number for the snRNA-seq data has been added to the Data Availability statement. A duplicated summary paragraph at the end of the Introduction has been removed.

## References

1. Raghu, G. et al. Idiopathic pulmonary fibrosis (an update) and Progressive pulmonary fibrosis in adults: An official ATS/ERS/JRS/ALAT clinical practice guideline. Am. J. Respir. Crit. Care Med. 205, e18–e47 (2022).

2. Ackermann, M. et al. Morphomolecular motifs of pulmonary neoangiogenesis in interstitial lung diseases. Eur. Respir. J. 55, 1900933 (2020).

3. Jacob, J. et al. Mortality prediction in idiopathic pulmonary fibrosis: evaluation of computer-based CT analysis with conventional severity measures. Eur. Respir. J. 49, 1601011 (2017).

4. Jacob, J. et al. Predicting outcomes in idiopathic pulmonary fibrosis using automated computed tomographic analysis. Am. J. Respir. Crit. Care Med. 198, 767–776 (2018).

5. Wollin, L. et al. Mode of action of nintedanib in the treatment of idiopathic pulmonary fibrosis. Eur. Respir. J. 45, 1434–1445 (2015).

6. Reininger, D. et al. Insights into the cellular and molecular mechanisms behind the antifibrotic effects of nerandomilast. Am. J. Respir. Cell Mol. Biol. 73, 700–712 (2025).

7. Richeldi, L. et al. Nerandomilast in patients with idiopathic pulmonary fibrosis. N. Engl. J. Med. 392, 2193–2202 (2025).

8. Ackermann, M. et al. The bronchial circulation in COVID-19 pneumonia. Am. J. Respir. Crit. Care Med. 205, 121– 125 (2022).

9. Murata, K. et al. Bronchial venous plexus and its communication with pulmonary circulation. Invest. Radiol. 21, 24–30 (1986).

10. Walsh, C. L. et al. Imaging intact human organs with local resolution of cellular structures using hierarchical phase-contrast tomography. Nat. Methods 18, 1532–1541 (2021).

11. Brunet, J. et al. Preparation of large biological samples for high-resolution, hierarchical, synchrotron phase-contrast tomography with multimodal imaging compatibility. Nat. Protoc. 18, 1441–1461 (2023).

12. Isensee, F., Jaeger, P. F., Kohl, S. A. A., Petersen, J. & Maier-Hein, K. H. nnU-Net: a self-configuring method for deep learning-based biomedical image segmentation. Nat. Methods 18, 203–211 (2021).

13. Hübner, R.-H. et al. Standardized quantification of pulmonary fibrosis in histological samples. Biotechniques 44, 507–511, 514–517 (2008).

14. Sikkema, L. et al. An integrated cell atlas of the lung in health and disease. Nat. Med. 29, 1563–1577 (2023).

15. Schupp, J. C. et al. Integrated single-cell atlas of endothelial cells of the human lung. Circulation 144, 286–302 (2021).

16. Adams, T. S. et al. Single-cell RNA-seq reveals ectopic and aberrant lung-resident cell populations in idiopathic pulmonary fibrosis. Sci. Adv. 6, eaba1983 (2020).

17. Engelbrecht, E. et al. Ectopic expansion of pulmonary vasculature in fibrotic lung disease and lung adenocarcinoma marked by proangiogenic COL15A1+ endothelial cells. Pulm. Circ. 15, e70102 (2025).

18. Leiber, L. M. et al. Aberrant and ectopic cell populations in restrictive allograft syndrome after lung transplantation. Eur. Respir. J. 2500537 (2026).

19. Newman, A. M. et al. Determining cell type abundance and expression from bulk tissues with digital cytometry. Nat. Biotechnol. 37, 773–782 (2019).

20. Steen, C. B., Liu, C. L., Alizadeh, A. A. & Newman, A. M. Profiling cell type abundance and expression in bulk tissues with CIBERSORTx. Methods Mol. Biol. 2117, 135–157 (2020).

21. Merritt, C. R. et al. Multiplex digital spatial profiling of proteins and RNA in fixed tissue. Nat. Biotechnol. 38, 586–599 (2020).

22. Gut, G., Herrmann, M. D. & Pelkmans, L. Multiplexed protein maps link subcellular organization to cellular states. Science 361, eaar7042 (2018).

23. Lang, N. J. et al. Ex vivo tissue perturbations coupled to single-cell RNA-seq reveal multilineage cell circuit dynamics in human lung fibrogenesis. Sci. Transl. Med. 15, eadh0908 (2023).

24. Raslan, A. A. et al. Lung injury-induced activated endothelial cell states persist in aging-associated progressive fibrosis. Nat. Commun. 15, 5449 (2024).

25. Skinnider, M. A. et al. Cell type prioritization in single-cell data. Nat. Biotechnol. 39, 30–34 (2021).

26. Truchi, M. et al. Aging affects reprogramming of pulmonary capillary endothelial cells after lung injury in male mice. Nat. Commun. 16, 7234 (2025).

27. Tan, Y. et al. SPACEc: a streamlined, interactive Python workflow for multiplexed image processing and analysis. Nat. Commun. 16, 10652 (2025).

28. Dimitrov, D. et al. LIANA+ provides an all-in-one framework for cell-cell communication inference. Nat. Cell Biol. 26, 1613–1622 (2024).

29. Mayr, C. H. et al. Spatial transcriptomic characterization of pathologic niches in IPF. Sci. Adv. 10, eadl5473 (2024).

30. Jin, S. et al. Inference and analysis of cell-cell communication using CellChat. Nat. Commun. 12, 1088 (2021).

31. Liebow, A. A. The bronchopulmonary venous collateral circulation with special reference to emphysema. Am. J. Pathol. 29, 251–289 (1953).

32. Turner-Warwick, M. Precapillary systemic-pulmonary anastomoses. Thorax 18, 225–237 (1963).

33. McDonough, J. E. et al. Transcriptional regulatory model of fibrosis progression in the human lung. JCI Insight 4, 131597 (2019).

34. Ackermann, M. & Konerding, M. A. Vascular casting for the study of vascular morphogenesis. Methods Mol. Biol. 1214, 49–66 (2015).

35. Ackermann, M., Mentzer, S. J., Kolb, M. & Jonigk, D. Inflammation and intussusceptive angiogenesis in COVID-19: everything in and out of flow. Eur. Respir. J. 56, 2003147 (2020).

36. Zheng, G. X. Y. et al. Massively parallel digital transcriptional profiling of single cells. Nat. Commun. 8, 14049 (2017).

37. Heaton, H. et al. Souporcell: robust clustering of single-cell RNA-seq data by genotype without reference genotypes. Nat. Methods 17, 615–620 (2020).

38. Danecek, P. et al. Twelve years of SAMtools and BCFtools. Gigascience 10, giab008 (2021).

39. Chang, C. C. et al. Second-generation PLINK: rising to the challenge of larger and richer datasets. Gigascience 4, 7 (2015).

40. Hoeft, K. et al. Label-free single-cell RNA multiplexing leveraging genetic variability. Nat. Commun. 15, 10612 (2024).

41. Young, M. D. & Behjati, S. SoupX removes ambient RNA contamination from droplet-based single-cell RNA sequencing data. Gigascience 9, giaa151 (2020).

42. Germain, P.-L., Lun, A., Garcia Meixide, C., Macnair, W. & Robinson, M. D. Doublet identification in single-cell sequencing data using scDblFinder. F1000Res. 10, 979 (2021).

43. McCarthy, D. J., Campbell, K. R., Lun, A. T. L. & Wills, Q. F. Scater: pre-processing, quality control, normalization and visualization of single-cell RNA-seq data in R. Bioinformatics 33, 1179–1186 (2017).

44. Hafemeister, C. & Satija, R. Normalization and variance stabilization of single-cell RNA-seq data using regularized negative binomial regression. Genome Biol. 20, 296 (2019).

45. Stuart, T. et al. Comprehensive integration of single-cell data. Cell 177, 1888–1902.e21 (2019).

46. Lopez, R., Regier, J., Cole, M. B., Jordan, M. I. & Yosef, N. Deep generative modeling for single-cell transcriptomics. Nat. Methods 15, 1053–1058 (2018).

47. Gayoso, A. et al. A Python library for probabilistic analysis of single-cell omics data. Nat. Biotechnol. 40, 163– 166 (2022).

48. Finak, G. et al. MAST: a flexible statistical framework for assessing transcriptional changes and characterizing heterogeneity in single-cell RNA sequencing data. Genome Biol. 16, 278 (2015).

49. Fang, Z., Liu, X. & Peltz, G. GSEApy: a comprehensive package for performing gene set enrichment analysis in Python. Bioinformatics 39, btac757 (2023).

50. Heumos, L. et al. Pertpy: an end-to-end framework for perturbation analysis. Nat. Methods 23, 350–359 (2026).

51. Schindelin, J. et al. Fiji: an open-source platform for biological-image analysis. Nat. Methods 9, 676–682 (2012).

52. Bankhead, P. et al. QuPath: Open source software for digital pathology image analysis. Sci. Rep. 7, 16878 (2017).

53. Palla, G. et al. Squidpy: a scalable framework for spatial omics analysis. Nat. Methods 19, 171–178 (2022).

54. Efremova, M., Vento-Tormo, M., Teichmann, S. A. & Vento-Tormo, R. CellPhoneDB: inferring cell-cell communication from combined expression of multi-subunit ligand-receptor complexes. Nat. Protoc. 15, 1484– 1506 (2020).

55. Hou, R., Denisenko, E., Ong, H. T., Ramilowski, J. A. & Forrest, A. R. R. Predicting cell-to-cell communication networks using NATMI. Nat. Commun. 11, 5011 (2020).

56. Zhou, Y. et al. Metascape provides a biologist-oriented resource for the analysis of systems-level datasets. Nat. Commun. 10, 1523 (2019).

57. van Buuren, S. & Groothuis-Oudshoorn, K. mice: Multivariate Imputation by Chained Equations in R. J. Stat. Softw. 45, 1–67 (2011).

